# Bioprinting of Human Primary and iPSC-derived Islets with Retained and Comparable Functionality

**DOI:** 10.1101/2024.10.14.617656

**Authors:** Miranda Poklar, K Ravikumar, Connor Wiegand, Ben Mizerak, Ruiqi Wang, Rodrigo M. Florentino, Zhenghao Liu, Alejandro Soto-Gutierrez, Prashant N. Kumta, Ipsita Banerjee

## Abstract

Currently, Type 1 diabetes (T1D) can be treated through implantation of allogenic islets, which replenish the beta cell population, however this method requires an extensive post-implantation immunosuppressant regimen. Personalized cellular therapy can address this through implantation of an autologous cell population, induced pluripotent stem cells (iPSCs). Cellular therapy, however, requires an encapsulation device for implantation, and so to achieve this uniformly with cells in a clinical setting, bioprinting is a useful option. Bioprinting is dependent on having a bioink that is printable, retains structural fidelity after printing, and is supportive of cell type and function. While bioprinting of pancreatic islets has been demonstrated previously, success in maintaining islet function post-printing has been varied. The objective of this study is to investigate the feasibility of printing functional islets by determining the appropriate combination of bioink, printing parameters, and cell configuration. Here, we detail the successful bioprinting of both primary human islets and iPSC-derived islets embedded in an alginate/methylcellulose bioink, with functionality sustained within the construct for both cell lineages. Sc-RNAseq analysis also revealed that printing did not adversely affect the genetic expression and metabolic functionality of the iPSC-derived islets. Importantly, the iPSC-derived islets displayed comparable functionality to the primary islets, indicating the potential to act as a cell source alternative for T1D implantation.

## 1. Introduction

In 2021, worldwide over 8.4 million individuals presented for Type 1 diabetes (T1D), with that number projected to escalate to 13.5-17.4 million cases by 2040. ^[4]^ In T1D, pancreatic β cells located in the islets of Langerhans are destroyed, leading to a severe insulin deficiency.^[5]^ Patients diagnosed with T1D are most commonly administered exogenous insulin to counteract this deficiency, however patients cannot always achieve glucose homeostasis, leading to cases of hypo- and hyperglycemia.^[6, 7]^ Current efforts are focused on restoring β cell mass and functionality within the patient through human cadaveric donor islet transplantation. In the last 20 years, these islets have been successfully implanted in human subjects, albeit the approach is heavily constrained by a lack of donor organs and an extensive immunosuppressant regimen required for the patient post-implantation.^[8, 9]^

Clinically, some progress has been made in developing biocompatible islet transplantation devices that can protect the transplanted islets from the host immune system. Transplanted islets are highly dependent on their physiological environment for growth and transportation of oxygen and nutrients, so most of the research has focused on developing hydrogel-based encapsulation devices. These macro-scale systems are designed to protect the cells in a structured 3D environment, while supporting and protecting the islets.^[10]^ Several of these devices are in early clinical stage, where embryonic stem-cell derived pancreatic endoderm cells (PEC-01) are macro-encapsulated in bioinert polymer capsules, and implanted in study patients for monitoring for the formation of an adequate β cell mass needed for functionality.^[11–13]^

Personalized medicine is a rapidly growing field with the desired tools to address these issues, where autologous cell therapies are utilized to avoid long-term allogenic rejection and the need for a continuous immunosuppressant drug regimen. One of the autologous cell therapies being discussed to alleviate the islet supply issue and the stringent immunosuppressant regimen needed is induced pluripotent stem cells (iPSCs). iPSCs have been successfully differentiated into islet mimetics (referred to here as iPSC-derived islets), and can act as a patient-specific cell source for islet transplantation.^[14–16]^ iPSC islets derived from the patients cells, not only have the potential to replace the β-cells destroyed from a T1D patient, they can also reduce the possibility of an allogenic immune rejection when implanted.^[17, 18]^ iPSCs additionally have the benefit of functioning as self-renewable single-source cells, therefore, demonstrating the potential to meet the scaleup of islet biomanufacturing necessary to satisfy the increased demand of islets for T1D patients. Beyond just the potential for transplantation, iPSCs are also excellent patient-specific cell sources for T1D disease modeling and drug discovery platforms.

While iPSCs are a promising foray into personalized cell therapy for the treatment of T1D, they still require a 3D micro-environment that can mimic the endocrine pancreas to function as intended. One technology that is designed to create this micro-environment is 3D bioprinting, which is an additive manufacturing technique that allows for the scale-up of islet organoid production. With iPSCs as the islet cell source, bioprinting can enable patient-specific cell therapy for T1D, by creating uniform and customizable scaffolds necessary for the function of the islets.^[19]^ In 3D bioprinting, specifically direct extrusion bioprinting, cells and bioink can be patterned and printed in a synergistic fashion using pressure as the driving force. These printed constructs can then closely mimic physiological soft tissue environments, allowing the transport of oxygen and molecules through the printed construct to the embedded cells. Bioprinting has thus over the years been successfully utilized for the printing of a variety of iPSC based soft tissue constructs, such as: cardiac, neural, and hepatic tissues, but attempts to print iPSC-derived pancreatic tissue have thus far yielded mixed results.^[20, 21]^

Given the importance of bioprinting islets, there have been considerable attempts at 3D printing islet constructs. Several studies have been able to show extended functionality when printing rat or porcine neonatal islet-like cell clusters (NICC) in an alginate/methylcellulose bioink, but these cell types are not analogous to human tissue for clinical purposes.^[22, 23]^ Parallel studies with printing of primary human islets and iPSC-derived islets have proved less encouraging. While overall there has been multiple reported success in printing human islets with retained viability, most of these printed islets failed to retain islet-specific function after printing. This could be attributed to the sensitivity of the islets to its microenvironment and to the stress incorporated during the process of 3D printing, which is likely to compromise islet function. Both primary and iPSC derived islets are highly sensitive to biophysical cues provided by their external surroundings, and any deviation from the appropriate mechanosignaling directives can result in stress and loss of function.^[24, 25]^ As documented in Table S1, the majority of islet printing studies where primary human islets were directly printed resulted in a loss of functionality, highlighting the sensitivity of islets to their environment.^[26]^ To date, there have also been no studies to the best of our knowledge involving the printing of functional iPSC-derived islets.

Motivated by the challenge of designing a tissue mimetic construct, our primary goal was to develop a robust technology to bioprint both primary human and iPSC-derived islets, preferably in a xeno-free bioink, with demonstrated viability and sustained GSIS function. This was accomplished by using a 3% w/v alginate/ 6% w/v methylcellulose bioink to encapsulate and print disparate trials of primary human islets, iPSC derived islets, and undifferentiated iPSCs in both, single cell, and aggregate form. Material characterization indicated that printing the bioink at pressures between 25-30 kPa and then crosslinking with calcium chloride produced macroporous structures that could facilitate adequate transportation of oxygen and nutrients, while maintaining a stiffness comparable to the pancreatic tissue. Both primary human islets and iPSC-derived islets were able to endure the stress of bioprinting without causing any significant effect on the viability of the cells. The bioprinted iPSC-derived islets and the primary human islets both successfully maintained functionality, indicating that with this bioink and printing protocol, diffusional limitations could be largely mitigated. Notably, the iPSC-derived islets exhibited similar levels of functionality to the printed primary human islets, which furthers adds validity to the idea that iPSC-derived islets are a promising alternative to primary human islets in future study. To the best of our knowledge, this is the first study that demonstrates extended functionality of both, primary human islets and iPSC-derived islets upon printing and extended culture. Finally, we were also able to show with single cell RNAseq conducted on bioprinted iPSC islets, that printing had no significant effect on the genetic expression or metabolic functionality of the iPSC islets in comparison to traditional culture methods. This serves to further solidify bioprinting as a valid alternative to traditional methods of maintaining iPSC islets for downstream analysis and personalized patient implementation.

## 2. Results

### 2.1. Alginate/Methylcellulose bioink provides the desired culture conditions for cell compatibility and structural support

A critical component in 3D bioprinting of islet constructs is to determine a bioink that retains a stable construct after printing, while supporting the viability and functionality of the printed islets. This bioink also needs to be printed under conditions that produce a high-resolution construct, where diffusional limitations potentially caused by the structure are mitigated. Achieving a successful printed construct is highly dependent on multiple parameters that affect the stability and viability of both the bioink and the embedded cells present, as shown in Figure 1a. Previous studies have reported printing viable human islets and functional rat islets with alginate/gelatin and alginate/methylcellulose bioinks respectively, however a bioink that can support both the extended viability and functionality of both primary and iPSC-derived islets has not yet been optimized.^[23, 27]^ Hence in this study we tested various combinations of alginate, methylcellulose and gelatin bioinks to determine the optimal bioink that could facilitate the extended culture of functional islets.

**Figure 1.**
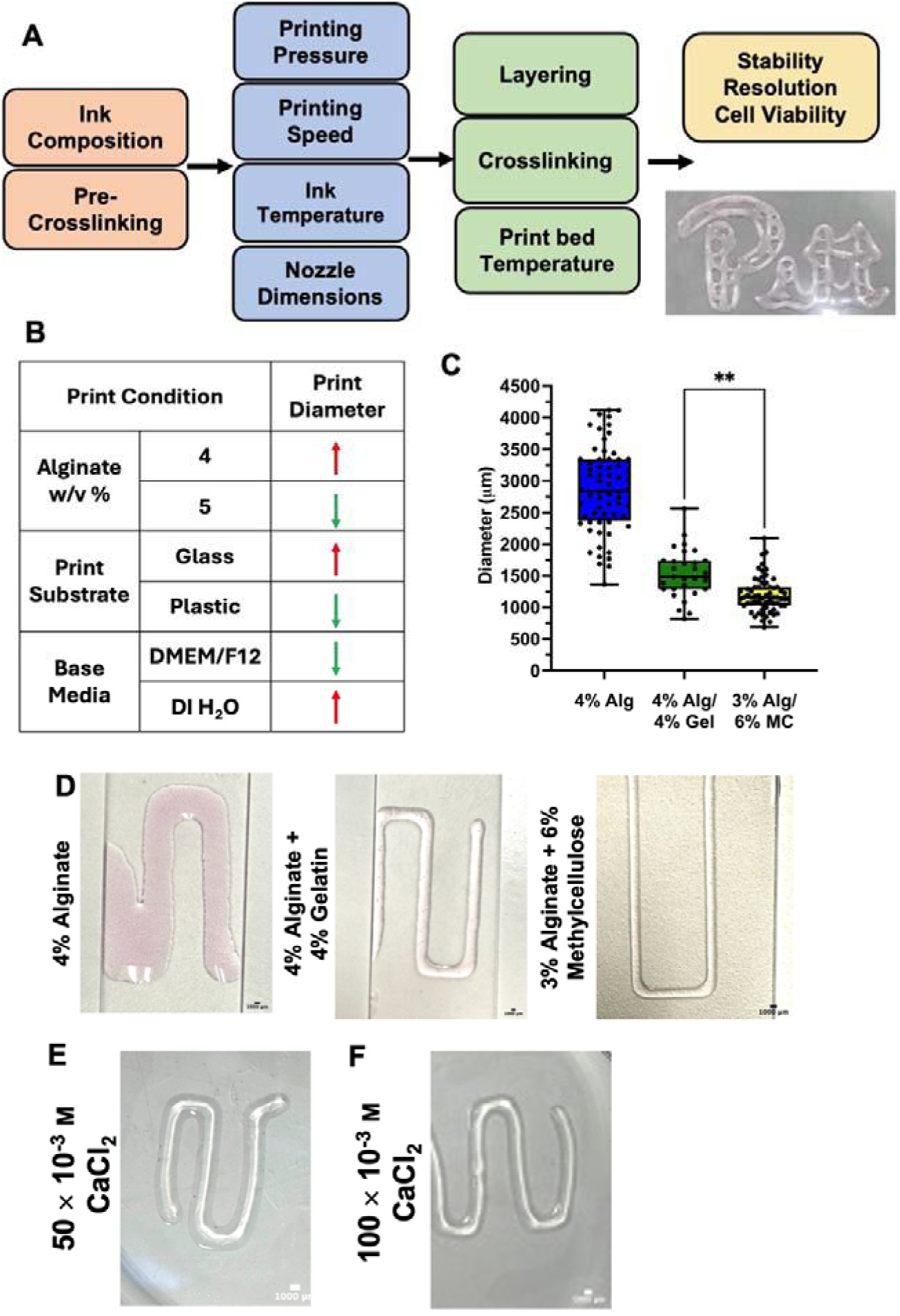
**A)** Schematic of independent parameters necessary for producing a stable and printed structure. **B)** Parameters that affect print diameter resolution in relation to each other. Red upward arrows indicate that the parameter causes the print diameter to increase in relation to the other parameter in the category. Green downward arrows indicate that the parameter causes the print diameter to decrease in relation to the other parameter in the category. All data can be found in Tables S2a-c. **C)** Diameter distribution for printed constructs comprised of either 4% w/v alginate, 4% w/v alginate/ 4% w/v gelatin, and 3% w/v alginate/ 6% w/v methylcellulose, printed using a 410 μm nozzle, at 15-45 mm s^-1^, and at pressures 8-30 kPa, pre-crosslinking. (**: p <0.01) **D)** Pre-crosslinked constructs composed of 4% w/v alginate, 4% w/v gelatin, and 3% w/v alginate/ 6% w/v methylcellulose, all printed using a 410 μm nozzle at 15 kPa and 30 mm s^-1^. **E)** 3% w/v alginate/ 6% w/v methylcellulose printed structure after 5 minutes of crosslinking using 50 × 10^-3^ M CaCl_2_. Printed at 30 kPa and 12 mm s^-1.^ **F)** 3% w/v alginate /6% w/v methylcellulose printed structure after 5 minutes of crosslinking using 100 × 10^-3^ M CaCl_2_. Printed at 30 kPa and 12 mm s^-1^.

To start optimizing the print diameter, alginate ink was printed at different concentrations, on different substrates, and formulated with different base media. It was seen that by increasing the alginate concentration, printing on a plastic substrate instead of glass, and formulating the ink with DMEM/F12 instead of DI water created higher resolution print structures (Figure 1b). All data formulating these conclusions can be found in Tables S2a-c in Supplemental Information. For each bioink a window of printability emerged, where only printing at a certain range of higher pressures and nozzle speeds could produce an even strand, rather than a series of droplets (Figure S1). When looking at the substrate to print on and the media that comprised the bioink, it was determined that the combination of printing on plastic substrates with an ink comprised of DMEM/F12 produced the highest resolution print construct.

One of the most consequential printing parameters that affects the resolution and fidelity of the construct is the composition of the bioink. To test the effect of bioink composition on the resolution of the printed construct, 4% w/v alginate, 4% w/v alginate/ 4% w/v gelatin (alg/gel), and 3% w/v alginate/ 6% w/v methylcellulose (alg/MC) were each printed at pressures between 8-30 kPa, and nozzle speeds between 15-45 mm s^-1^ onto plastic substrates. The alg/gel bioink also required extra heating of the ink before extrusion in order to prevent formation of a granular hydrogel, which requires a higher printing pressure and reduces the print resolution.^[28]^ The alginate and alg/gel bioinks produced structures that had an average diameter of 2861 ± 665 μm and 1532 ± 388 μm, respectively (Figure 1c). In the case of the alg/MC bioink, when printed at the same pressures and nozzle speeds, constructs were on average 1198 ± 273 μm in diameter. The difference in printing resolution between the bioinks was also qualitatively evident, as the alginate and alg/ gel bioinks exhibited spreading after printing at 15 kPa and 30 mm s^-1^, in comparison to the alg/MC ink (Figure 1d).

Having determined that alg/MC bioink produces the highest resolution printed construct in comparison to other bioinks tested, this bioink was next examined for its crosslinking capabilities. Ionic crosslinking of the structures increases the print resolution and the construct becomes structurally intact.^[29, 30]^ The 3% alg/ 6% MC bioink was crosslinked with either 50 × 10^-3^ M CaCl_2_ or 100 × 10^-3^ M CaCl_2_, for 3, 5 or 10 minutes (Table S2d). The ink when crosslinked with 50 × 10^-3^ M CaCl_2_ displayed no obvious deformities when crosslinked for 5 minutes (Figure 1e) but was not fully formed when crosslinked for 3 minutes (Figure S2). Crosslinking with 100 × 10^-3^ M CaCl_2_ resulted in deformities in the printed construct, with non-uniform swelling all throughout, regardless of the crosslinking time (Figure 1f). Rheological characterization of the printed construct was accomplished using atomic force microscopy (AFM) micro-indentation with a colloidal probe.^[31]^ AFM in liquid contact mode utilizing force-volume measurements (Figure S3 a, b) on the printed constructs produced a Young’s modulus of 7.86 ± 3.71 kPa and a stiffness of 11.59 ± 4.16 mN m^-1^ (Figure 2a). A preliminary surface topography map was also determined using the colloidal probe (Figure S3c).

**Figure 2.**
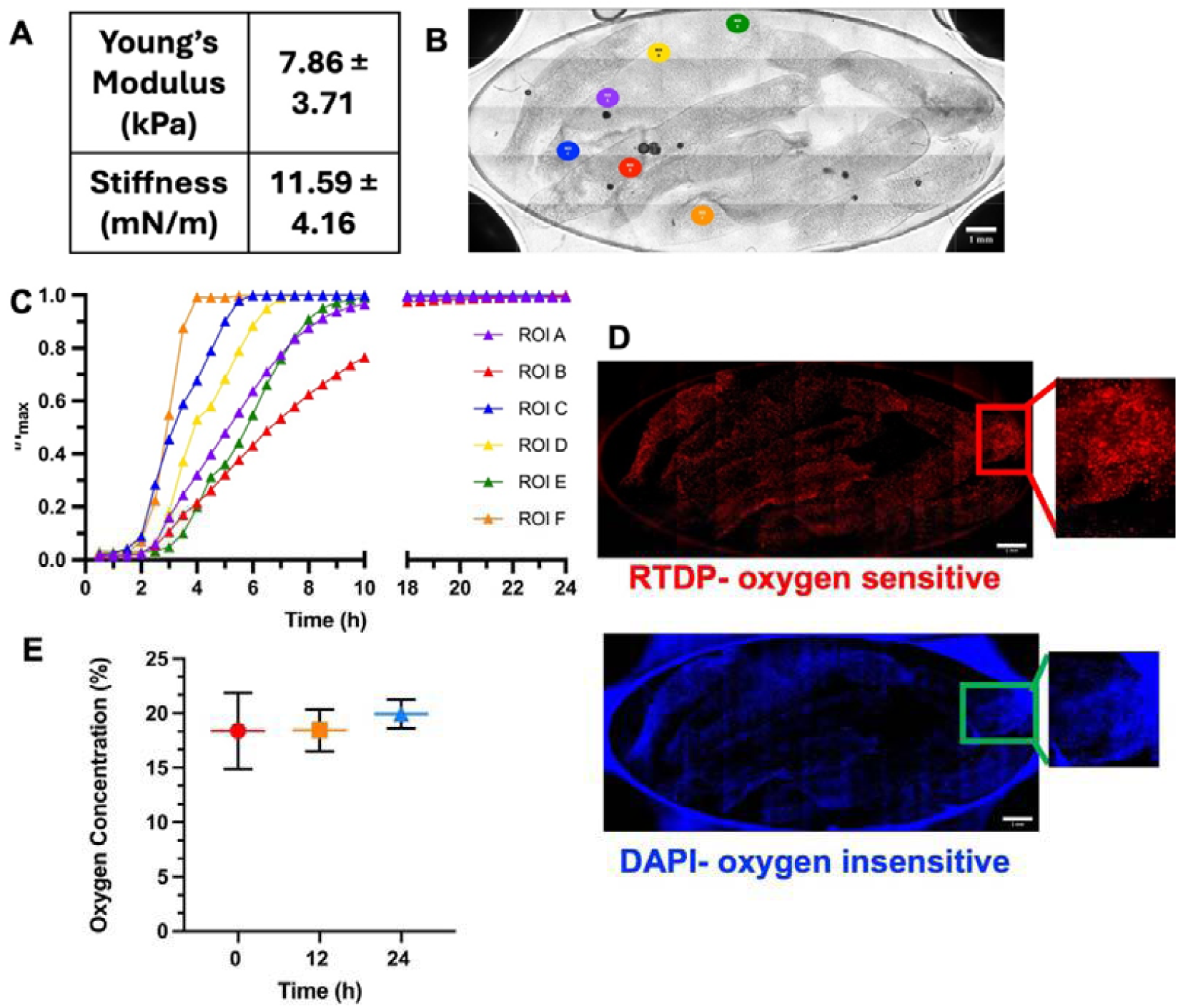
**A)** Young’s modulus (kPa) and stiffness (mN m^-1^) of printed construct crosslinked with 50 × 10^-3^ m CaCl_2_ measured by AFM force-volume contact mode in liquid. **B)** Phase imaging of the printed structures placed on the middle layer membrane of the Micronit device, with ROI placement indicated. **C)** Fluorescent intensity ratio of specific ROIs under FITC-Dextran (70 kDa) flow for 24 hours. A break in the X axis was included from hours 10-18. **D)** Printed structures containing embedded RTDP beads (oxygen sensitive) and DAPI (oxygen insensitive beads), which were loaded on a Micronit chip and fluorescently imaged over the course of 24

Beyond the print resolution and structural stability, the printed construct needs to provide adequate oxygen diffusion for cell survival, and additionally, facilitate the transport of glucose to and insulin from the embedded islets. To quantify the potential of the printed construct to efficiently transport oxygen and nutrients, it was placed on a microfluidic device membrane and monitored under constant FITC-dextran (70 kDa) flow for 24 hours. The FITC-dextran was used to track molecular diffusion within the print through measuring the fluorescence intensity. When measured at specific regions of interest (ROIs) (Figure 2b) a clear increase of intensity with time was noted, indicating diffusion of 70 kDa dextran into the construct, albeit with some variations in the dynamics of diffusion based on the location of the ROI in the flow field (Figure 2c). The initial ‘dead time’, where fluorescence intensity did not change with respect to time, was observed to be ∼2 hours, as this is the time required to attain the detectable FITC concentration in the system. All 6 of the ROI’s reached steady state, where the change in fluorescent intensity with respect to time is constant, within 1.5 hours after ‘dead time’. From steady state at each of the ROIs, the average diffusion coefficient for 70 kDa dextran through the printed construct was estimated to be 7.14 x 10^-6^ cm^2^ s^-1^. Based on these conclusions, it is a reasonable assumption that 0.180 kDa glucose and 5.8 kDa insulin molecules would be able to successfully diffuse in and out of the printed construct within an appropriate time frame when loaded with islets.^[32]^

In parallel, the printed structures were also loaded with RTDP and DAPI beads to report the oxygen concentration within the printed construct (Figure 2d). The RTDP bead fluorescence was quenched in the presence of oxygen, and the DAPI beads acted as a control, providing an intensity ratio that was proportional to oxygen concentration in the system (Figure S4). Representative images of bead fluorescence at 0% and 18% oxygen can be found in Supplemental Information, Figure S5. Over the course of 24 hours, the printed constructs maintained an oxygen concentration of ∼18%, indicating that the printed structure retained ambient oxygen concentration throughout the duration of the experiment (Figure 2e).

### 2.2 Alginate/methylcellulose bioink preserves phenotype and function of bioprinted human islets

After establishing that the alg/MC ink could support proper nutrient and oxygen diffusion throughout the printed construct, the next progression was to test the feasibility of printing primary human islets in the designed bioink. Primary human islets are difficult to culture and maintain *in-vitro*; however, given an appropriate microenvironment they can exhibit robust *in-vitro* function.^[33–35]^

Human cadaveric primary islets were printed in the alginate/methylcellulose ink and maintained in culture for 3 to 7 days (Figure 3a). Details regarding the donor can be found in Supplemental Information (Table S3). 500 islets were handpicked and mixed into 0.5 mL bioink using a syringe LuerLock system. The constructs were printed at 25 kPa and a nozzle speed of 12 mm s^-1^, crosslinked for 5 minutes in 50 × 10^-3^ M CaCl_2_, and cultured in PIM(R) media. Live Dead imaging (Figure 3b) 3 days after printing demonstrated that the printed islets retained viability and their characteristic spheroidal morphology required to maintain islet function. There was no significant difference noted in structure between printed primary human islets, and non-printed control primary human islets (Figure S6).On average, 93.8 ± 2.6 % of the total area of each islet (Figure 3c) was considered live, with most of the dead cells being located closer to the periphery of the islet (Figure 3d). The printed islets were next tested for phenotype by immunostaining for Glucagon and C-peptide, which are key hormones produced by the α and β cells respectively. C-peptide is analogous to insulin, as it is attached to insulin in the proinsulin form and cleaved when insulin is released from β cells.^[36]^ As shown in Figure 3e, printed islets had appropriate cytosolic expression of C-peptide and Glucagon, indicating the presence of both α and β cells dispersed throughout the islet.

**Figure 3.**
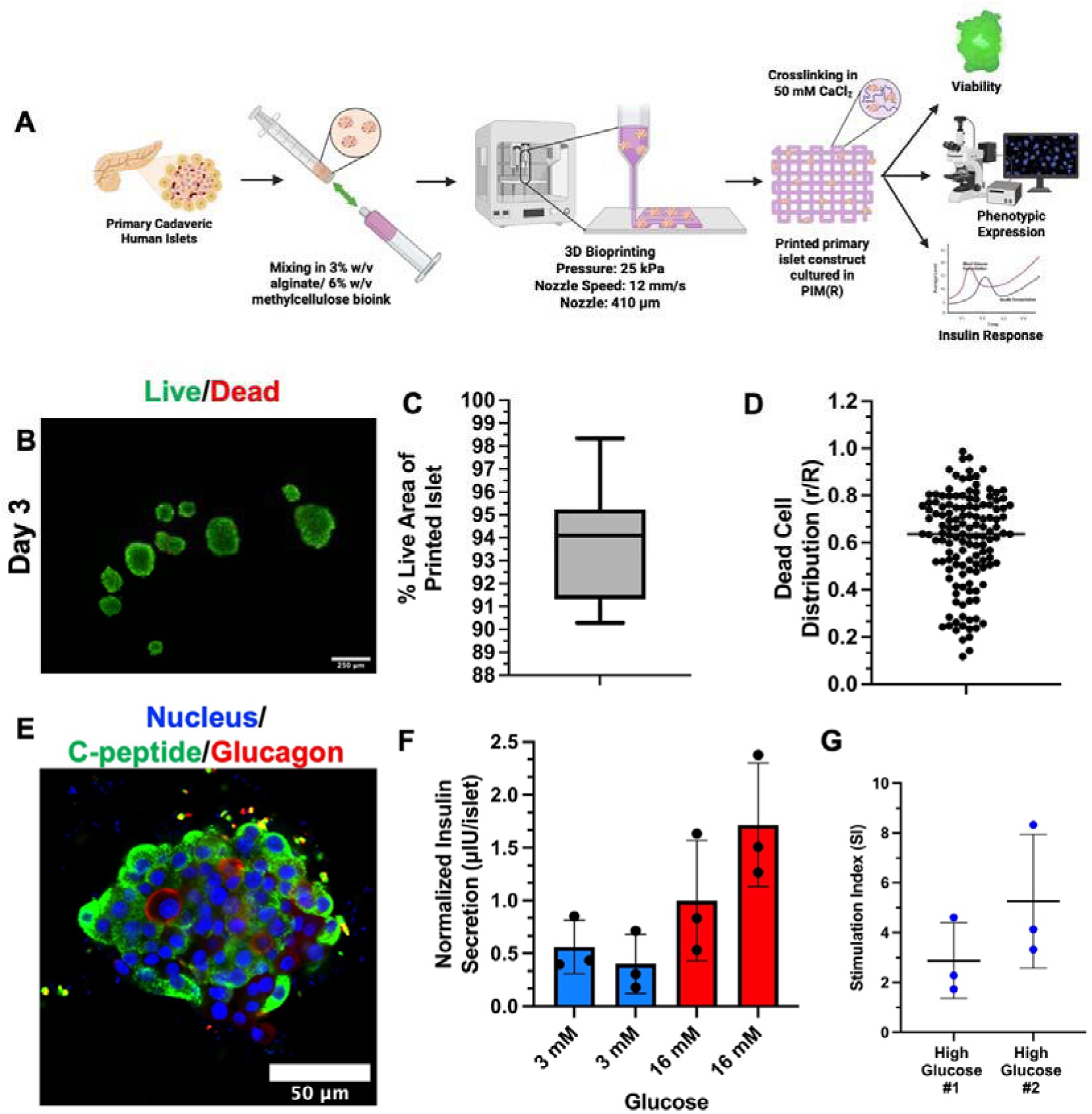
**A)** Schematic view of printing primary cadaveric human islets embedded in a 3% w/v alg/ 6% w/v MC bioink. The printed primary islet construct is then cultured for 3-7 days to monitor for functionality and viability. Created using Biorender.com.^[2]^ **B)** Live Dead staining of printed primary islets 3 days post-printing. All scale bars convey 250 μm. Live cells are shown in green, dead cells are shown in red. **C)** Percentage of individual printed primary islet represented by viable cells, as indicated by green signal in Live/Dead staining (N=8). **D)** Location of dead cells present in individual printed primary islet as indicated by red signal in Live Dead staining. r/R of 0.0 indicates that the dead cell is present in the center of the islet, and an r/R of 1.0 indicates that the dead cell is present at the edge of the islet. **E)** Primary islets were printed and maintained in culture for 7 days. Islets are fluorescently stained for nuclei (DAPI), islet marker C-peptide (shown in green), and islet marker Glucagon (shown in red). Scale bar representing 50 μm (*: p < 0.05). **F)** Glucose stimulated insulin secretion of printed primary islets after 7 days in culture **G)** Stimulation index from GSIS of first high glucose period normalized to second low glucose period (n=3).

Finally, the function of the bioprinted islets were tested after 7 days of culture post-printing. Islet function is measured by its ability to secrete controlled amount of insulin in response to glucose, termed Glucose Stimulated Insulin Secretion (GSIS). Islet GSIS function is tested *in-vitro* by measuring the secreted levels of insulin in response to consecutive exposure to low (3 × 10^-3^ M) and high glucose (16 × 10^-3^ M) conditions. The insulin secretion of control, non-printed primary human islets can be found in Supplemental Information: Figure S7. In the current set-up the islets are residing inside the bioprinted constructs and could encounter a diffusion barrier, which can skew the observed GSIS response. To test for the transport limitation, we modified the GSIS protocol by testing for 2 sequential low and high glucose exposure. As shown in Figure 3f, there was no significant difference between the two sequential low and concurrently the two sequential high glucose exposure, indicating adequate transport of glucose and insulin across the construct. Further, there was a significant increase in insulin secretion when exposed to the first round of high glucose (average of 0.99 ± 0.57 μΙU insulin per islet) after the low glucose condition (an average of 0.40 ± 0.28 μΙU insulin per islet) resulting in an impressive stimulation index (SI, first round of high glucose/second round of low glucose) of 2.87 ± 1.52 (Figure 3g).

### 2.3 Single-cell iPSCs demonstrate the ability to form aggregates within printed constructs

Having demonstrated the feasibility of bioprinting human islets in the designed alginate/methylcellulose bioink, our next objective was to test this ink for printing iPSCs and iPSC-derived islets. We first tested with printing iPSCs in their pluripotent stage, followed by inducing differentiation post-printing.

Adherent cultured iPSC-2s were first dissociated into single cells and mixed at a cell seeding density of 1.6 million cells per mL of alginate/methylcellulose bioink. Cells were printed and maintained for 8 days in propagating media and monitored for aggregate formation (Figure 4a), which is advantageous for islet differentiation. ^[37, 38]^. As shown in Figure 4b, cells started aggregating over time, with distinct aggregates being visible by Day 4 of culture. Phase imaging taken at 0, 2, 4, and 8 days post-printing indicated that once aggregated, it maintains its spheroidal structure while growing in size comparable to an average islet of Langerhans, between 100-200 μm, around Day 8 (Figure 4c).^[39]^ Live Dead staining of the aggregates within the printed construct indicated substantial dead cell presence (Figure 4d); however harvesting the aggregates by dissolving the bioink with EDTA show high viability of the individual aggregates and insignificant red signal (Figure 4e). This indicates that the dead cells in the printed construct were likely debris and non-aggregated cells in the strand. However, once aggregated they are stable with minimal cell death within the aggregate. Thus, it is feasible to print iPSCs as single cell suspension in the bioink and culture them into stable aggregates post-printing; however, it should be noted that this process results in significant amount of debris in the printed construct. Assuming the dead cells to be single cells that failed to aggregate, one approach to eliminate cell death is to pre-form iPSC aggregates, followed by printing of aggregates, which is what we attempted next.

**Figure 4.**
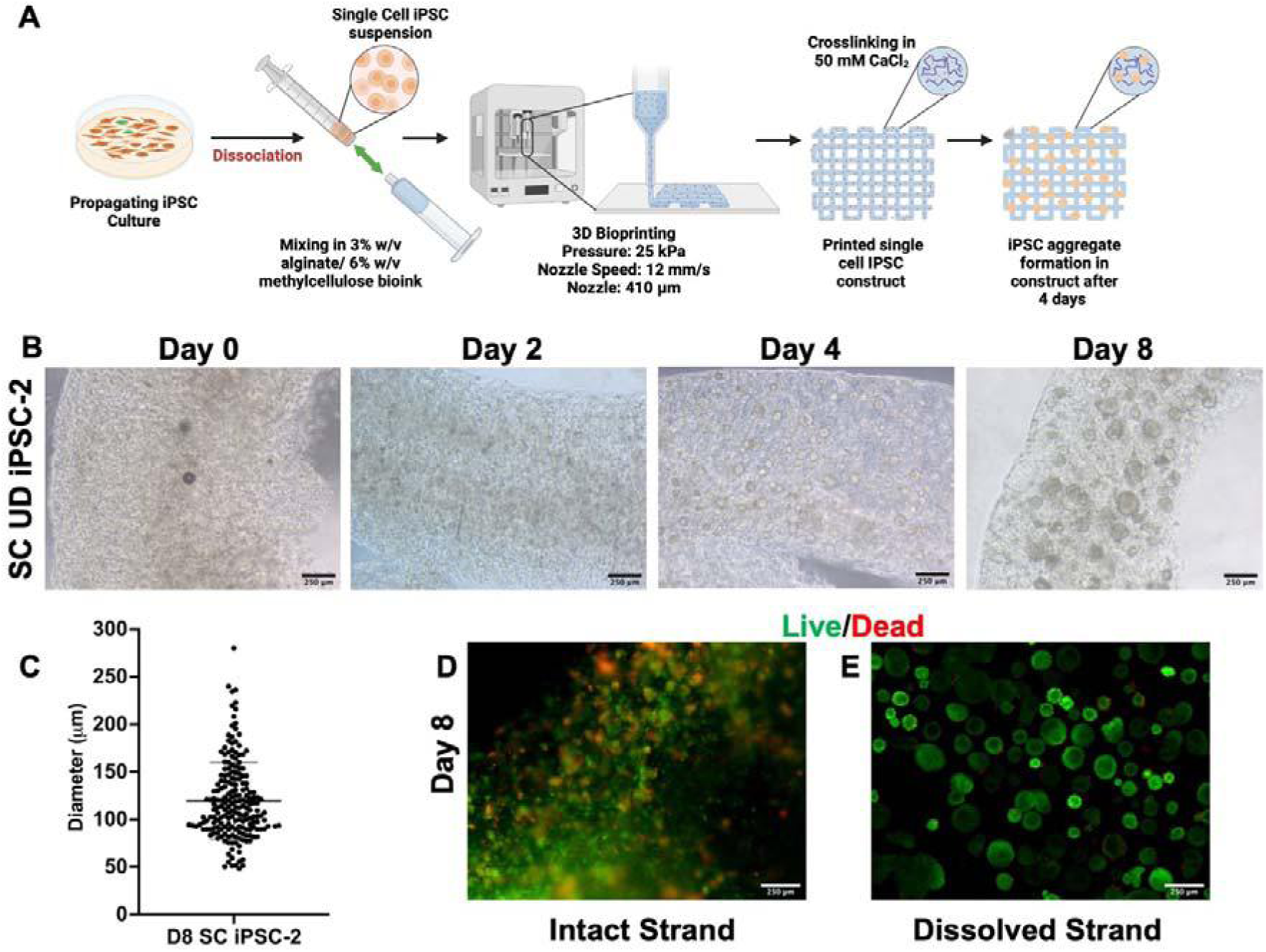
**A)** Schematic view of printing single cell iPSCs embedded in a 3% w/v alg/ 6% w/v MC bioink. The printed single cell construct is then cultured for 8 days to monitor aggregate formation. Created using Biorender.com.^[1]^ **B)** Phase images (4x), taken 0, 2, 4, and 8 days after printing demonstrating aggregate formation within the strand. Scale bars represent 250 μm. **C)** Live Dead staining of printed single cell iPSC-2s within the printed structure 8 days after printing. All scale bars convey 250 μm. Live cells are shown in green, dead cells are shown in red. **D)** Live Dead staining conducted after the printed structure was dissolved, showing single cell iPSC-2 UD aggregate formation after 8 days cultured within the printed structure. **E)** Size (diameter) distribution of aggregates formed after 8 days of culture in the strand. Aggregates were decapsulated from strand and imaged using phase microscopy.

### 2.4 3D printed iPSC aggregates maintain pluripotency and demonstrate potential for successful differentiation within the printed construct

Our group has previously reported the fabrication of microwells to facilitate uniform iPSC aggregation.^[40]^ As detailed in the materials and methods, propagating iPSCs were dissociated into single cells upon confluency and seeded on the agarose microwells, where they aggregated overnight, generating aggregates 250 μm in diameter.

Two different in-house generated iPSC cell lines, iPSC-1 and iPSC-2 were aggregated, and 3D printed following the previously detailed protocol and cultured in the printed construct for 3 days in a mTeSR1 media with 10 × 10^-6^ M Y-27632 (Figure 5a). To compare the effect of 3D printing on the iPSC aggregates, control aggregates (iPSC-aggregates that have not gone through the printing process) were taken from the same microwells, manually encapsulated in 1.5% w/v alginate and cultured for the same duration and using the same media as the printed cells. The control condition was chosen to maintain similar culture configuration and biophysical cues as the 3D printed cells.

**Figure 5.**
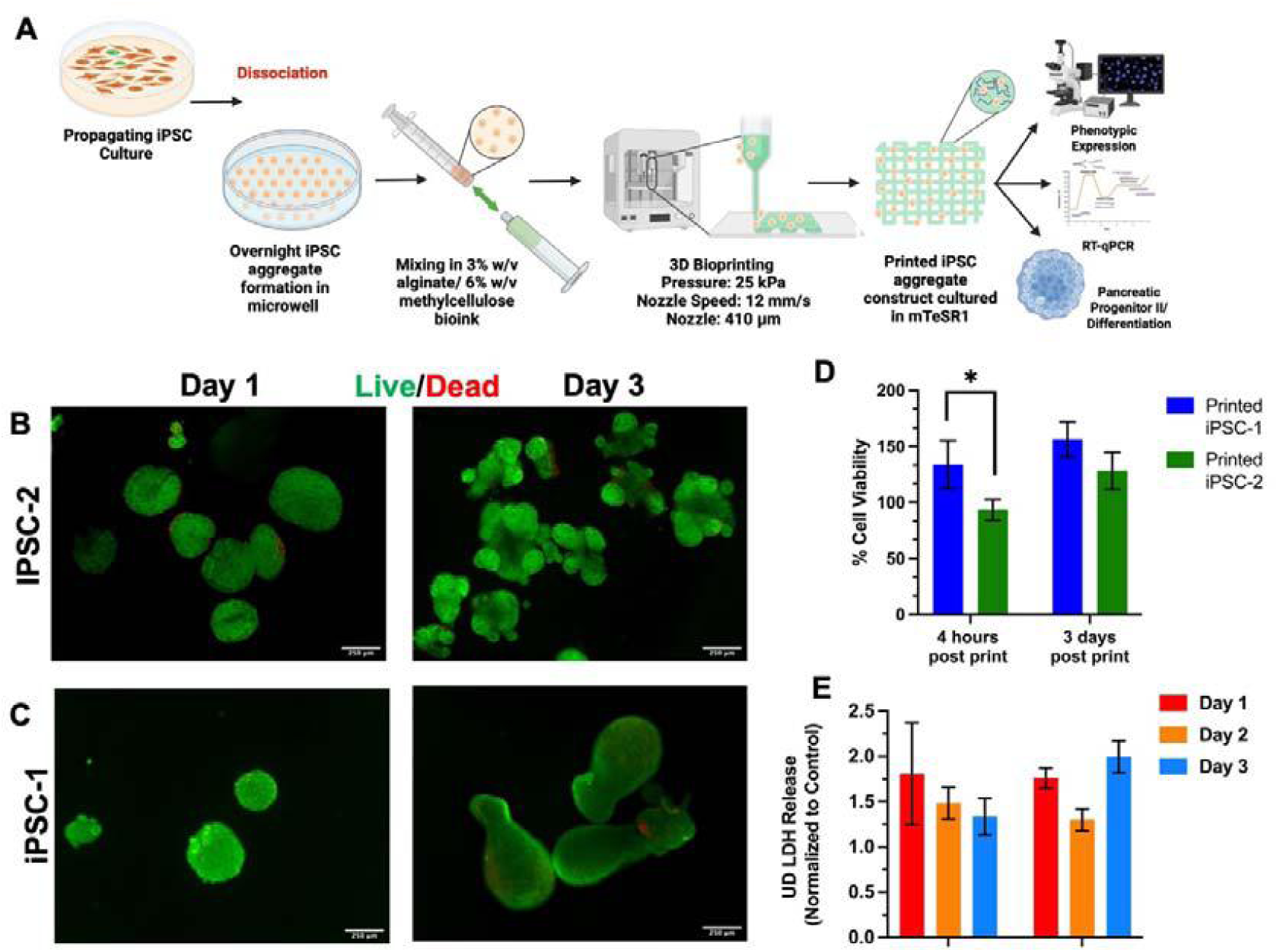
**A)** Schematic view of printing aggregate IPSCs embedded in a 3% w/v alg/ 6% w/v MC bioink. Propagating cells are dissociated upon confluency, added to microwells and incubated overnight for aggregate formation. Aggregates are printed and maintained in culture for further analysis. Created using Biorender.com.^[3]^ **B)** Live Dead staining of printed iPSC-2 UD aggregates 1- and 3-days post-printing. All scale bars convey 250 μm. Live cells are shown in green, dead cells are shown in red. **C)** Live Dead staining of printed iPSC-1 UD aggregates 1- and 3-days post-printing. All scale bars convey 250 μm. Live cells are shown in green, dead cells are shown in red. **D)** Undifferentiated iPSC-2 and iPSC-1 aggregates from individual printed constructs stained for MTS 4 hours and 3 days after printing. Printed aggregates were normalized with control aggregates encapsulated in 1.5% w/v alginate. N = 3 (*: p < 0.05). **E)** Media from iPSC-2 and iPSC-1 UD prints (N=3) and iPSC-2 and iPSC-1 UD controls was collected for 3 days and tested for LDH release. Printed aggregates were normalized with control aggregates encapsulated in 1.5% w/v alginate.

Live Dead staining on Days 1 and 3 post printing, showed high aggregate viability and minimal cell death for both the cell lines (Figure 5 b, c). Interestingly, by Day 3, it was evident that the iPSC aggregates were coalescing within the printed construct, resulting in an increase in aggregate size by an average of 184% for iPSC-1, and 129% for the iPSC-2 over Day 1 aggregates (Figure S9). The viability of both the cell lines, as quantified using MTS assay (Figure 5d) was high even after 3 days of culture, and cell death, as quantified by LDH release (Figure 5e), remained consistent between the controls and both printed cell lines. Thus overall, the printing procedure and bioink supported viability of both the cell lines with minimal cell death.

Next, the printed aggregates were tested for pluripotency after 3 days of culture, by immunofluorescent staining for key pluripotency markers, OCT4 and NANOG. Both the iPSC cell lines exhibited overall excellent expression of NANOG and OCT4 (Figure 6a, b). In both the cases, however, a spatial variation of the stain was noted, with the stain being rather weak or absent towards the core of larger aggregates. The gene expression level of *OCT4* and *NANOG* was also verified through RT-qPCR, where the printed aggregates expression levels were normalized to control aggregates. Both the tested iPSC cell lines exhibited similar or an increase in gene expression compared to control (iPSC-1: 0.78 for OCT4, 1.21 for *NANOG*; iPSC-2: 1.65 for *OCT4* and 1.87 for *NANOG* (Figure 6c)), demonstrating retained pluripotency after printing. Comparing across the two iPSC lines, iPSC-2 line had significantly higher levels of gene expression for pluripotency markers (*: p < 0.05) over iPSC-1. Thus, all future experimentation was continued on with the iPSC-2 line.

**Figure 6.**
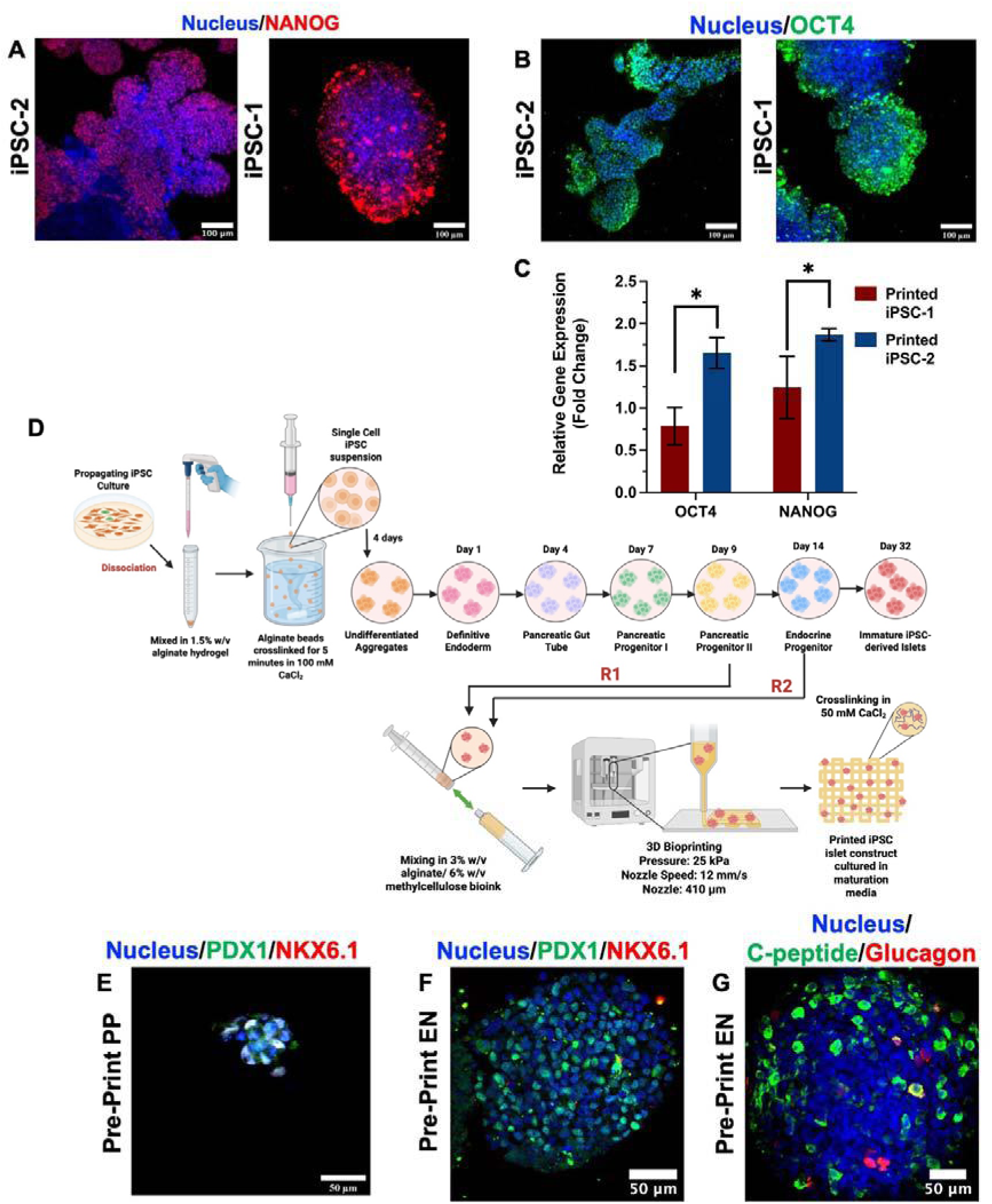
**A)** iPSC-2 and iPSC-1 undifferentiated iPSC aggregates fluorescently stained after 3 days for nuclei (DAPI), and pluripotency marker NANOG, shown in red. Scale bar depicts 100 μm. **B)** Undifferentiated iPSC-2 and iPSC-1 iPSC aggregates stained after 3 days in culture conditions. Aggregates are fluorescently stained for nuclei (DAPI), and pluripotency marker OCT4, shown in green. Scale bar representing 100 μm. **C)** RT-qPCR quantification of ΔΔC_t_ fold change in genes of interest *NANOG* and *OCT4* in undifferentiated iPSC-2 and iPSC-1 aggregates after 3 days cultured in printed strand. The level of the gene expression is normalized to the control aggregates encapsulated in 1.5% w/v alginate. Error bars represent the standard deviation from three independent printed structures (*: p < 0.05). **D)** Schematic view of iPSC islet differentiation, starting with an adherent iPSC culture which is dissociated and mixed at a ratio of 1 million cells: 3 mL 1.5% w/v alginate. The cell-alginate mixture is dropped into 100 × 10^-3^ M CaCl_2_ and crosslinked, creating cell loaded alginate beads which expand into aggregate loaded beads over 4 days. The iPSC aggregates are then differentiated into 6 different developmental stages, culminating in a mature iPSC islet form. R1 denotes aggregates printed at the pancreatic progenitor (PP) II stage, and R2 denotes aggregates printed at the endocrine progenitor (EN) stage. Created with BioRender.com.^[44]^ **E)** Pancreatic progenitor (PP) aggregates were decapsulated after 2 weeks in differentiation culture conditions, and then printed and maintained in culture for 3 days. Aggregates are fluorescently stained for nuclei (DAPI), pancreatic progenitor marker PDX1 (shown in green), and pancreatic progenitor marker NKX6.1 (shown in red). Scale bar representing 50 μm. **F)** Endocrine progenitor aggregates were decapsulated after 24 days in differentiation culture conditions, and then printed and maintained in culture for 3 days. Aggregates are fluorescently stained for nuclei (DAPI), pancreatic progenitor marker PDX1 (shown in green), and pancreatic progenitor marker NKX6.1 (shown in red). Scale bar representing 50 μm. **G)** iPSC-2 aggregates that differentiated to the endocrine progenitor stage (EN) and printed were also fluorescently stained for islet markers C-peptide (shown in red), and Glucagon (shown in green). Scale bar representing 50 μm.

The next milestone was to induce differentiation of the printed pluripotent aggregates through different stages of islet development. Accordingly, differentiation was induced over 32+ days following the chemical agents in a previously reported directed differentiation protocol.^[16, 41, 42]^ Over the differentiation period, the aggregates within the printed strand appeared to lose their structure and their spherical morphology, and produced a significant amount of debris, as seen in Figure S10. Upon decapsulation, the aggregates fragmented and failed to maintain their structure. Attempts to differentiate the aggregates to an earlier developmental stage, pancreatic progenitor II (14 days of culture) showed mixed results. The aggregates exhibited similar patterns of increasing amounts of debris and fragmentation throughout differentiation (Figure S11). However, the aggregates could still be immunostained and imaged, where the aggregates expressed key markers expected at the pancreatic progenitor II stage, PDX1 and NKX6.1 (Figure S12).

We thus aborted the plan to print in the pluripotent stage and instead considered pre- differentiating prior to printing. The pre-differentiation was conducted following our previously reported procedure of encapsulating IPSCs in 1.5% w/v alginate capsules and differentiating within the alginate capsule to the subsequent developmental stage (Figure 6d).^[43]^ The aggregates thus differentiated to specific stages were retrieved, suspended in the bioink, and printed using the protocol previously detailed. Both the iPSC pancreatic progenitor II aggregates (Figure 6d, R1) and the endocrine progenitor aggregates (Figure 6d, R2) retained integrity during the printing and maintained their morphology and viability 3 days after printing (Figure S13, S14). Importantly, both groups of printed aggregates displayed key developmental markers, PDX1 and NKX6.1, after 3 days in print culture (Figure 6e, f respectively). The printed endocrine progenitor aggregates also expressed C-peptide and Glucagon after printing (Figure 6g), both markers that indicate the transition of the aggregates to insulin-producing cells.

### 2.5 Printing of iPSC-derived islets maintained their phenotype and functionality post printing

With more success in printing pre-differentiated cells, we next attempted to further differentiate the aggregates to the stage of immature islets prior to bioprinting (Figure 6d). The iPSC islets were decapsulated from the alginate beads, suspended in the 3% w/v alg/ 6% w/v MC bioink at an average density of 500 aggregates per 0.5 mL of bioink, printed at 25 kPa and a nozzle speed of 12 mm s^-1^, and then cultured with maturation media for 7 days (Figure 7a).

**Figure 7.**
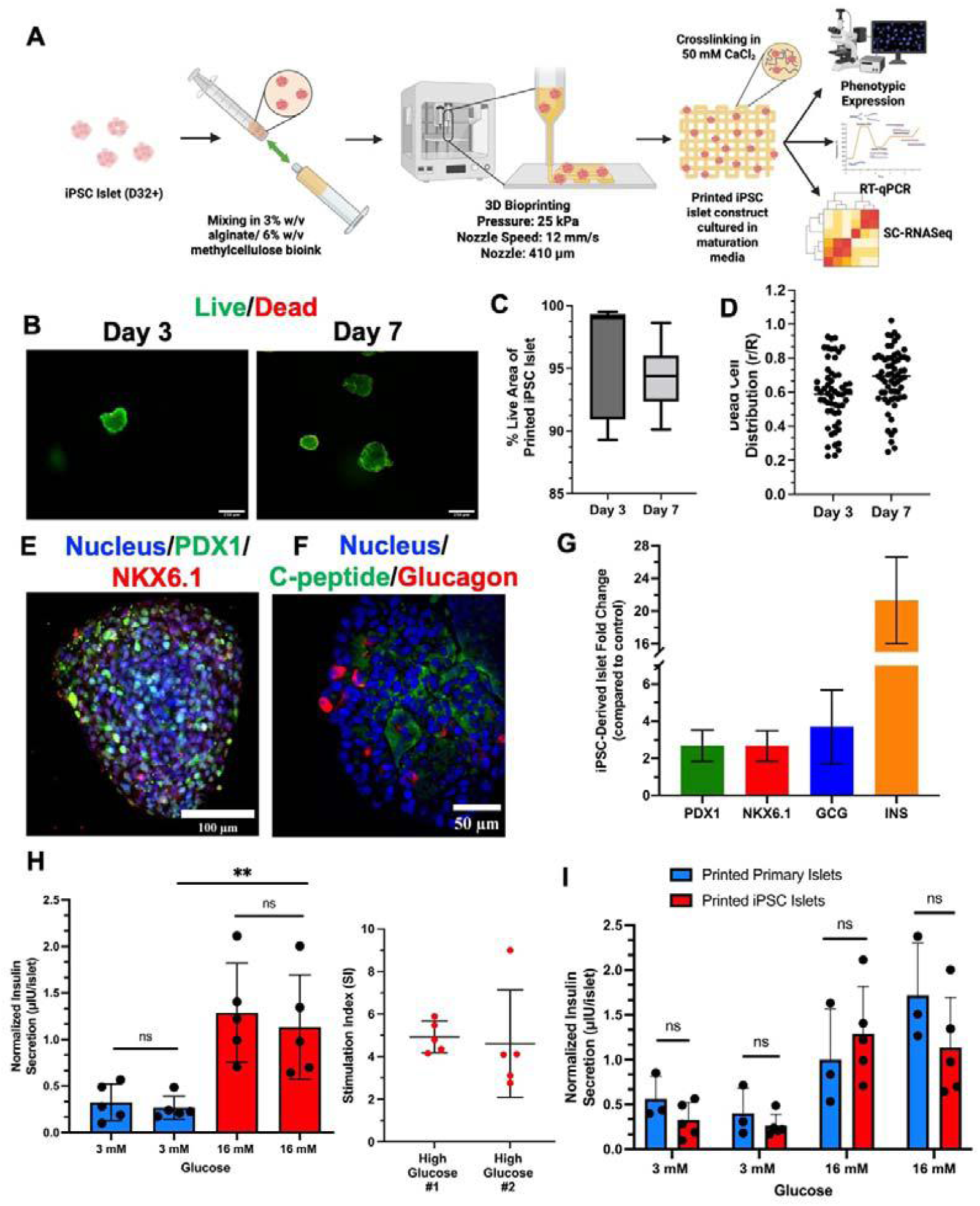
**A)** Schematic view of printing iPSC-derived islets embedded in a 3% w/v alg/ 6% w/v MC bioink. The printed construct is kept in culture for 3-7 days with maturation media. Created with Biorender.com.^[45]^ **B)** Live Dead staining of printed iPSC-derived islets 3- and 7-days post-printing. All scale bars convey 250 μm. Live cells are shown in green, dead cells are shown in red. **C)** Percentage of individual printed iPSC islets represented by viable cells, as indicated by green signal in Live Dead staining (Day 3: n =5, Day 7: n=10). **D)** Location of dead cells present in individual printed IPSC islets as indicated by red signal in Live Dead staining. r/R of 0.0 indicates that the dead cell is present in the center of the islet, and an r/R of 1.0 indicates that the dead cell is present at the edge of the islet. **E)** iPSC-derived islets were decapsulated after differentiation and then printed and maintained in culture for 7 days. Aggregates are fluorescently stained for nuclei (DAPI), pancreatic progenitor marker PDX1 (shown in green), and pancreatic progenitor marker NKX6.1 (shown in red). Scale bar representing 100 μm. **F)** Aggregates are fluorescently stained for nuclei (DAPI), islet marker C-peptide (shown in green), and islet marker Glucagon (shown in red) after 7 days of culture. Scale bar representing 100 μm. **G)** RT-qPCR quantification of ΔΔC_t_ fold change in genes of interest *PDX1*, *NKX6.1*, Glucagon (*GCG*), and Insulin (*INS*) in printed mature iPSC-derived islets and Y1 aggregates after 7 days cultured in printed strand. The level of the gene expression is normalized to the control immature iPSC islets encapsulated in 1.5% w/v alginate. Error bars represent the standard deviation (n= 3). **H)** Glucose stimulated insulin secretion of printed iPSC islets after 7 days in culture. Stimulation index normalized to second low glucose period (n=5) (*: p < 0.05, **: p < 0.01, ***: p < 0.005, ****: p < 0.001). **I)** Combined GSIS of printed primary (n = 3) and iPSC islets (n = 5), (*: p < 0.05, **: p < 0.01).

Live Dead staining performed 3 and 7 days after printing showed that the iPSC-derived islets retained their structural integrity when decapsulated from the construct (Figure 7b). It was also noted that no significant amount of debris was present within an intact construct before decapsulation, as seen in Figure S15. In comparison to control iPSC-derived islets, the morphology remained similar, however in the printed cells, small sections of cell outgrowth were noted (Figure S16). Quantification of Figure 7b revealed high post-print viability of the islet organoids, with 95.9% and 94.3% (Figure 7c) viability at post-printing days 3 and 7 respectively. Further quantification of the spatial location of the few dead cells that were present in the organoids revealed more of a presence towards the periphery of the islet (Figure 7d). The printed organoids were further characterized for their gene and protein expression; immunofluorescent staining for key markers showed appropriate co-expression of NKX6.1 and PDX1 (Figure 7e), and expression of C-peptide and Glucagon (Figure 7f), with no co-expression of those present.

RT-qPCR was also conducted in order to examine the gene expression of *PDX1*, *NKX6.1*, *GCG*, and *INS*. These gene expression levels were compared to control iPSC islets, which were unprinted islets that had undergone one less week of maturation in comparison to the printed iPSC islets. For pancreatic progenitor genes *PDX1* and *NKX6.1*, the printed iPSC islets demonstrated log_2_ fold changes of 2.68 ± 0.84 and 2.67 ± 0.83, respectively (Figure 7g). In the case of *GCG*, a log_2_ fold change of 3.70 ± 1.99 was observed (Figure 7g). Most importantly, for *INS*, a key marker for islet maturation, the printed islets exhibited a fold change of 21.30 ± 5.30 (Figure 7g), in comparison to the control iPSC islets. This is a favorable indication that the printed islets can mature within the printed construct, and display the appropriate gene expression.^[46]^ and thus were next tested for functionality.

7 days post-printing GSIS was conducted, with the printed iPSC islets being exposed to two rounds of low glucose (3 × 10^-3^ M), and two rounds of high glucose (16 × 10^-3^ M), following an identical experimental protocol to what was used in the testing of the printed primary islets. The insulin secretion of control, non-printed iPSC-derived islets can be found in Supplemental Information: Figure S17. The printed iPSC islets exhibited excellent functionality, with the insulin secretion at the second low glucose period averaging to 0.26 ± 0.13 μΙU insulin per islet, and that of the first high glucose period averaging to 1.29 ± 0.53 μΙU insulin per islet(Figure 7h), resulting in an SI (first high glucose period/ second low glucose period) of 4.93 ± 0.74 (Figure 7h). What was most notable however, was that there was no significant difference in insulin secretion levels between the printed primary human islets and the printed iPSC islets at any incubation period, low or high (Figure 7i). These results indicate that iPSC islets can be successfully printed, mature within the printed construct, and maintain comparable functionality levels to primary human islets while in extended culture.

### 2.6 scRNA-seq Analysis indicates that printed iPSC-derived islets do not experience significant metabolic stress

Islets are very sensitive to changes in their micro-environment, and hence, to better understand how the stress of printing and culture in the printed constructs may affect the iPSC-derived islets, a comprehensive gene expression study was conducted using single-cell RNA sequencing. Thus, 7 days after printing, the iPSC islets were harvested and sequenced in comparison to iPSC control islets from the same batch, cultured for the same duration.

From sequencing a list of differentially expressed genes (DEG) between the printed islets and the control islets was identified. This DEG list was further filtered to only include statistically significant genes that had >1.0 or <-1.0 Log2Ratio gene expression. Ingenuity Pathway Analysis (IPA) was utilized to determine the biologically significant canonical pathways that were altered with printing, with 25 relevant pathways (p < 0.05) demonstrated in Figure 8. Several key pathways related to islet functionality were upregulated in the printed iPSC islets, including insulin secretion signaling (z-score = 7.247, p < 0.005), calcium signaling (z-score= 6.143, p < 0.0001), and gap junction signaling (z-score = N/A, p < 0.001), which drives intra-islet beta cell signaling.^[47]^ When focusing on pathways involved in cell stress, such as NRF2 oxidative stress response signaling and endoplasmic reticulum (ER) stress response, none of the aforementioned pathways showed any statistically significant upregulation or downregulation in contrast to control islets. Autophagy (z-score = 3.628, p < 0.0005) however is upregulated in the printed sample, and has been shown to act as a protectant against ER stress.^[48]^ Conversely however, the Type II diabetes mellitus pathway is upregulated (z-score: 3.55 with a bias, p < 0.0001) and the PPARα/ RXRα pathway (z-score = -1.987, p < 0.001) is downregulated in comparison to the control iPSC islets, indicating the possibility of some islet dysfunctionality within the printed samples. Further examination, however, of the endocrine cell specific population revealed that significant differences in gene expression level were only noted in 1/7 genes of interest in the ER and oxidative stress pathways (Figure S18). This suggests that the islets were not significantly adversely affected in terms of cellular stress in comparison to the control iPSC islets, thus demonstrating the efficacy of the bioprinting process.

**Figure 8.**
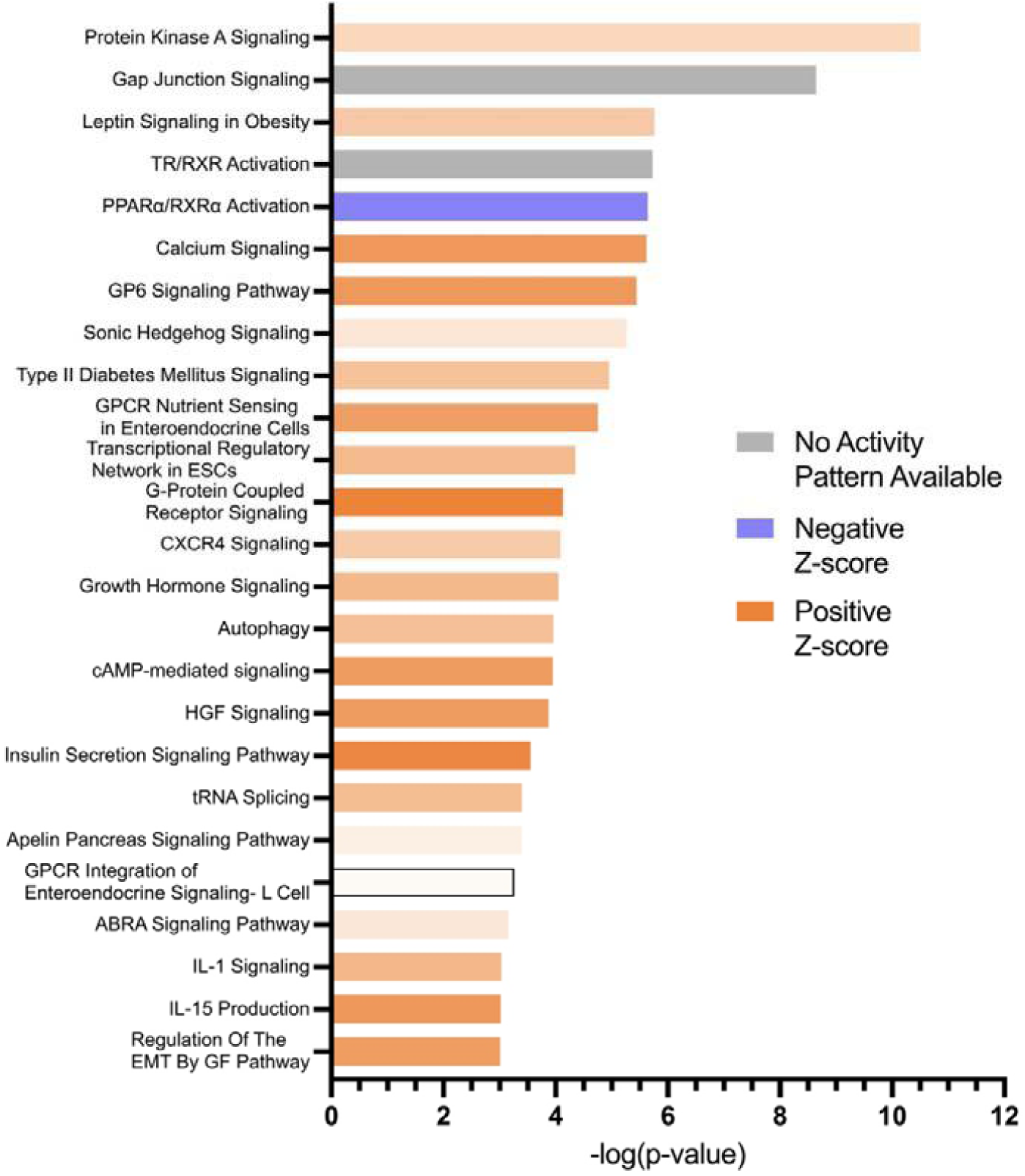
Ingenuity Pathway Analysis (IPA) for 25 biologically relevant significantly altered (p <0.05) canonical pathways for the printed construct in comparison to the control. Z score is displayed as a gradient color scale, with a darker blue or orange indicating a higher Z score. A blue Z score denotes a downregulated pathway, an orange Z score denotes an upregulated pathway, and a gray Z score indicates that an activity pattern for that pathway could not be established.

## 3. Discussion

In this work we demonstrated direct extrusion 3D bioprinting with an alginate/methylcellulose bioink to be a viable and promising option for the printing and extended culture of functional primary and iPSC-derived islets. The printed constructs formed after ionic crosslinking of the bioink can provide the appropriate environment necessary for the iPSC islets to maintain their phenotype and functionality; very encouragingly, the iPSC islet function was found to be comparable to that of printed primary human islets. By printing iPSC-derived islets, we can thus, provide a scaffold necessary for further exploration into patient-specific islet transplantation. We were also able to demonstrate by scRNAseq analysis that none of the major stress pathways are being upregulated from the printing.

A key aspect of cellular bioprinting is to converge on an optimal bioink which retains structural fidelity while also supporting cell viability and function, immediately after printing and in the long run. The immediate effect of printing on the cells are primarily governed by printing parameters, while long term effects are dictated by the cell microenvironment, influenced by the physio-chemical properties of the ink. However, these parameters cannot be decoupled and implemented individually, they are all simultaneously at play and need to be determined as such.

Alginate based bioinks have thus far, been the most preferred and commonly used bioinks for printing pancreatic islets and relevant cell population. In our group we have used alginate extensively to encapsulate iPSCs and derive functional pancreatic islet like cells^[43, 49–51]^. Marchioli et. al. successfully printed viable primary human islets in a 4% w/v alginate and 5% w/v gelatin bioink, with the caveat that these printed islets start to lose functionality one day after printing. This was theorized to be caused by diffusional limitations of the alginate/gelatin bioink, as the islets recovered functionality after decapsulation on Day 7.^[27]^ Another study had also included alginate as the bioink basis when printing mouse islets, but also saw reduced functionality more than one day after printing.^[52]^

Duin et. al, were able to address this issue in their functional rat and NICC islet studies, by including methylcellulose in their bioink (3% w/v alginate/ 9% w/v methylcellulose).^[22, 23]^ Both porcine neo-natal islet-like cell clusters (NICC) or rat islets maintained functionality and viability; 7 days in the case of the rat islets, and 28 days for NICC.^[22, 23]^. It was hypothesized that because the methylcellulose eluted out of the construct within 3 days of printing, a microporous structure was created, resulting in unimpeded nutrient and oxygen diffusion.^[53]^ Even with this functional success however, there still exists a gap in transferring this progress to clinically applicable cells.

While alginate based bioinks are more predominantly used for islet printing, Clua-Ferré et. al saw sustained functionality when rat pancreatic β-cell INS1E cells encapsulated in collagen-tannic acid (1% w/v tannic acid) were printed into spheroids.^[54]^ The other class of natural bioink is one derived from the native tissue to maximize cellular compatibility. Hwang et. al. used pancreatic decellularized ECM (pdECM) hydrogel as a bioink to print H1 embryonic stem cell (ESC) derived endocrine progenitor cells, and were able to demonstrate that printed aggregates were able to maintain viability and respond appropriately to glucose.^[55]^

In our work, primary human islets that were printed in the 3% w/v alg/ 6% w/v MC bioink retained function after 7 days of culture; GSIS showed that the islets responded appropriately to low and high glucose conditions, resulting in an average SI of 2.87 ± 1.52 (first high glucose incubation/second low glucose incubation). Importantly, the GSIS was conducted on the intact printed construct, meaning that no diffusional limitations were present that could significantly hinder the functionality of the primary islet, even in extended culture. The lack of a diffusional barrier was also verified through the inclusion of two rounds of low glucose exposures followed by two rounds of high glucose exposure. If an increase in secretion was noted between the two low glucose exposures or the two high glucose exposures, this delay in insulin secretion could be indicative of a diffusion barrier. In both the primary and iPSC-derived islet cases, no significant difference was noted between the two low exposures or two high exposures. It is worth noting that several groups had previously shown that primary human islets could remain viable when implanted in 3D bioprinted structures, however in most of these studies the islets themselves were not printed but incorporated in a printed scaffold. ^[33, 56, 57]^

While alginate has been a widely adopted biocompatible polymer for cell encapsulation and drug delivery purposes, the combination of methylcellulose and alginate offers many advantages. A previous study printing mesenchymal stem cells in 3% w/v alg/ 9% w/v MC and crosslinked with 100 × 10^-3^ M CaCl_2_ demonstrated that the inclusion of methylcellulose creates a macroporous print structure, which is not present in an alginate only bioink.^[53]^ The addition of methylcellulose serves to enhance the porosity of the structure in extended culture, as the methylcellulose is not crosslinked, and therefore elutes out. Methylcellulose is also considered to be biocompatible, and has been shown not to induce a severe foreign body response or necrosis even after 8 weeks in vivo.^[58]^ While the methylcellulose in the printed constructs does leave and reduce the stability of the structure, the crosslinked alginate can still be manipulated for potential implantation, as evidenced by a video taken four days after printing (Movie S1). While in our study we did not image the bioink ultrastructure, we did measure the potential effect of the bioink on the embedded cells, in terms of the diffusional properties and structural attributes. When placed under continuous perfusion for 24 hours fluorescently tagged dextran (70 kDa), comparable in size to glucose and insulin, was able to penetrate even to the middle of the printed 3% alginate/ 6% methylcellulose construct within 1 hour. In addition, it was estimated that the diffusion coefficient of the 70 kDa dextran in the construct was 7.14 x 10^-6^ cm^2^ s^-1^. While there are no direct comparisons found in literature, Leddy et. al. found that the diffusion coefficient for 70 kDa dextran in 2% w/v alginate was 1.5 x 10^-6^ cm^2^ s^-1^, indicating that our diffusion coefficient is an appropriate order of magnitude.^[59]^

The phenotype and function of the cells, in particular stem cells, are well known to be sensitive to the mechanical and physical constitution of the tissue micro-environment.^[60–62]^, in this case due to the biophysical and biochemical nature of the ink. It is expected that reproducing the native tissue microenvironment will be most suitable for in-vitro tissue culture^[63–67]^. Rubiano et. al. reported the steady state modulus of resected pancreatic tissue to be 1.06 ± 0.25 kPa, and in the case of rat pancreatic tissue, our lab has reported a Young’s modulus of 1.21 ± 0.077 kPa.^[68, 69]^. Testing the mechanical properties of the printed alg/MC construct with AFM we obtained an average Young’s modulus of 7.86 kPa. Thus, the elasticity of our printed construct is comparable in magnitude to that of a native pancreatic environment, albeit a bit higher. We attribute the Young’s modulus of the printed construct to the use of both a low viscosity alginate and calcium chloride as the crosslinker.

As mentioned earlier, cell viability during bioprinting process is largely dependent on the printing conditions, including nozzle diameter, and printing pressure; these same parameters also govern the resolution of the print, but with the inverse relationship. Furthermore, the print resolution will influence the necessary diffusion of oxygen and nutrients to and from the embedded cells. Hence there is significant nonlinear interdependence between parameters and conditions, highlighting the optimal requirement for an ideal bioink as being one that can produce high resolution scaffold at low printing pressures and speeds, such that transport limitations are minimized, thereby preserving viability of the printed cells.^[70]^ Klak et. al found that when printing primary human islets with a 3% w/v alginate bioink and a 600 μm nozzle, the optimal printing pressure needed to be maintained below 30 kPa as printing higher than this significantly impacted the viability of the islets, due to the increase in shear stress.^[71]^. In our study we tested multiple alginate-based bioinks, of which the 3% w/v alg/ 6% w/v MC bioink produced the highest resolution structures when printed at pressures below 30 kPa and nozzle speeds between 5-45 mm s^-1^. Each construct was printed as a 3D single layer, with a focus on optimizing the bioink to facilitate the maintenance of phenotype and function. While in this paper we are not focused on multilayer structures, the feasibility of 3D printing in multiple layers has been demonstrated and can be implemented with the current system when needed.^[72–74]^

Beyond printing, the physicochemical property of the construct is also dictated by the method and concentration of crosslinking, which directly affects the cell fate. For alginate structures, ionic crosslinking is primarily employed, as alginate can bind readily with multivalent cations. The 3% w/v alg/ 6% w/v MC bioink was crosslinked with 50 × 10^-3^ M CaCl_2_ for 5 minutes, and a stable structure was produced with no deformations present. It is possible however, to utilize other divalent cations such as barium and strontium, and/or explore higher crosslinking concentration while keeping the printing parameters constant. ^[30]^ While this would have increased the print resolution, thereby reducing diffusion barrier, it would have also increased the bioink stiffness modulus hence negatively affecting the cell fate.

In printing iPSC-derived tissues we need to account for the dynamics of differentiation as an additional printing parameter, since the requirement for microenvironmental niche is specific to the stage of organ maturation. Therefore, we evaluated printing the iPSCs at different stages of differentiation, starting from the pluripotent stage. Our first attempt was to print the iPSCs in the initial propagation stage, as single cells suspended in the bioink, and continue with the process of cell aggregation and subsequent differentiation within the printed construct. When single cell iPSCs were printed at a cell density of 1.6 million cells mL^-1^ of alg/MC bioink, aggregate formation was noted by Day 4 of culture, however a significant amount of cell debris was observed. The aggregates that formed were uniform, spherical, viable, and remained intact after decapsulation, indicating that even with the debris present, aggregation was successful. Similar observation was reported by Li et. al when printing single cell iPSCs in a hydroxypropyl chitin bioink, with a cell density of 1 million cells mL^-1^ bioink, which resulted in significant amount of dead cell debris. ^[75]^ Uniform aggregation of the single cells was also observed and aggregation in the more porous and softer bioink (lower hydroxypropyl chitin concentration) was attributed to a combination of both migration and in situ proliferation. Conversely, Gu et. al, were able to print single cell iPSCs at a much higher density of 40 million cells per 0.5 mL of an agarose/carboxymethyl-chitosan/alginate bioink with minimal cell death and debris present.^[76]^ While this study indicates the feasibility of minimizing single cells by increasing cell seeding density, we took the route of pre-aggregating the iPSCs prior to printing instead of printing single cells.

Printing of iPSC aggregates were tested at different stages of differentiation, which revealed interesting stage-specific dynamics, likely arising from the differential balance of proliferation and differentiation. The printed iPSC aggregates in both lines coalesced and formed larger aggregates within the construct, but clearly maintained pluripotency and viability, demonstrating the potential for induction towards a desired lineage by specific chemical cues. Coalescing was also clearly observed with differentiation post-printing to the pancreatic progenitor or endocrine progenitor stage within the printed construct, where the aggregates not only merged, but also produced a significant amount of debris around the aggregate. While these aggregates displayed phenotypic expression of PDX1 and NKX6.1 they were not structurally stable and fragmented upon decapsulation. This phenomenon, however, was significantly reduced by pre-differentiating the aggregates prior to printing, most likely from the suppression of proliferation with differentiation.

The iPSC-islets printed at the pre-mature stage followed by post-printing maturation were able to maintain structure and viability over 7 days during which they displayed a significant increase in insulin gene expression in comparison to control iPSC islets, indicating the printed environment as being supportive of maturation. In the same manner as the printed primary human islets, GSIS conducted after 7 days indicated that the iPSC-islets could appropriately respond to glucose, resulting in an average SI of 4.93 ± 0.74. What was very promising was that in comparison to the insulin secretion exhibited by the printed primary human islets under the same GSIS conditions, there was no significant difference in insulin secretion levels between the two populations, demonstrating that the iPSCs were comparatively functional. To the best of our knowledge this is the first work demonstrating bioprinting of both primary and iPSC-islets with preserved post-printing function. A summary of previously reported primary human islets and iPSC-derived islets is included in the Supplemental Information Table S1, with a focus on the duration of culture and reported functionality. It is important to note that while there has been some previous success in reporting post-printing viability of the islets, many of these studies failed to maintain islet functionality after printing. In contrast to those reports, our study successfully retained islet function for both primary human islets and iPSC-derived islets when tested a week after printing.

It is still relatively unknown the extent of stress the cells tend to undergo when printed, therefore, we conducted scRNASeq to determine the alterations made to key stress and islet specific metabolic pathways in comparison to control iPSC islets that has not undergone printing. As seen in Figure 8, key islet pathways such as calcium signaling and insulin secretion signaling are significantly upregulated in comparison to the control iPSC islets, supporting the results of the functionality and phenotypic assays conducted on the printed islets. In comparison to the control, stress pathways ER stress, and NRF-2 oxidative stress mediated response were not significantly altered. ER stress in islet cells can be induced when demand for insulin protein folding overcomes the capabilities of the β cells, and with oxidative stress the cells are not able to keep up with an increase in reactive oxygen species.^[77]^ For cells specifically within the endocrine cell population, it was determined that the expression levels of only 1/7 genes of interest from both stress pathways were significantly different, highlighting that printing is not causing undue stress on the islets.

Although the insulin secretion signaling pathway of the printed iPSC islets was significantly enriched in comparison to the control iPSC islets and there was no clear activation of metabolic stress pathways, there are indicators that printing and culturing of iPSC islets in the printed constructs may in some ways adversely affect key metabolic pathways involved in insulin secretion. As seen in Figure 8, PPARα/ RXRα activation was significantly downregulated (z-score = -1.987, p < 0.001) in the printed iPSC islets in comparison to the control islets. Activation of the PPARα/ RXRα pathway is vital for regulating fatty acid metabolism, and it has been shown that in PPARα/ RXRα knockout mice insulin secretion is inhibited.^[78, 79]^ It was also shown that the Type II diabetes mellitus pathway is significantly enriched (z-score: 3.55, p < 0.0001). Salg et. al. conducted sc-RNAseq transcriptome analysis on printed INS-1 cells, and hypothesized that stress induced by bioprinting, hyperglycemia, and hypoxic conditions may lead to overexpression of the IGF-1/TGFβ signaling pathways and consequent insulin-producing cell inflammation.^[80]^

## 4. Conclusion

This project expands the possibilities for high-throughput patient-specific treatment of T1D diabetes, by successfully combining the advantages of bioprinting and iPSCs. iPSC-derived islets and primary islets were able to withstand the rigors of printing and remain functional even after extended culture in the printed construct. Most notably, the printed iPSC islets displayed similar levels of functionality to the printed primary islets, strengthening the idea that iPSC-derived islets have the potential to replace primary islets as the main source of T1D implantation treatment. This printing technology also shows the promise for printing differentiated tissues other than islets. Further development of this technology could explore the potential for in-vivo implantation of printed iPSC-derived islets for not only T1D treatment, but also for in-vitro patient-specific drug testing and disease modeling.

## 5. Materials and Methods

### Primary Islet Isolation and Culture

Human cadaveric pancreatic islets were procured from Prodo Labs (San Francisco, USA) and maintained in suspension using the proprietary Prodo islet media for recovery (PIM(R)) with media changes every 2-3 days, which maintains a fasting islet state at 5 × 10^-3^ M glucose. The details of the donor can be found in Supplementary Information (Table S3).

Islets were maintained in a 100 mm × 25 mm petri dish (Fisher Scientific) suspended in PIM(R) (7-10 mL). The islets were initially cultured for 7 days in PIM(R) under static suspension conditions to recover from isolation and transport. A 200 μL pipette with SureOne aerosol barrier pipette tips (Fisher Scientific) was used to transfer individual islets for mixing in the bioink with the aid of a dissecting microscope.

### iPSC Culture and Encapsulation

All iPSCs in these experiments were progeny of two iPSC cell lines, Y1, denoted iPSC-1, and hFL262 iPSC, denoted iPSC-2. The details of generation and characterization of iPSC-2 can be found in the Supplemental Materials and Methods. Undifferentiated iPSCs were encapsulated as a single cell suspension according to our previous study.^[43]^ Further details describing cell dissociation and encapsulation can be found in Supplemental Materials and Methods.

### iPSC Differentiation into Pancreatic Islet Endocrine Cells

Using a modified version of a previously published protocol islet-like cells were generated using the stagewise differentiation steps as described in Supplemental Material and Methods. ^[16, 41, 42]^

### Alginate/Gelatin and Alginate/ Methylcellulose Bioink Preparation

For all the bioinks that contained an alginate component, alginic acid sodium salt derived from brown algae was used (A2158, MilliporeSigma). To prepare the alginate bioink, 4% w/v alginate was dissolved in DMEM/F12 (Gibco) at 35 °C for one hour. This protocol was also followed when developing 1.5% w/v alginate. The 4% alginate/ 4% gelatin bioink was synthesized by stirring 4% w/v alginate into DMEM/F12 at 35 °C for one hour. 4% w/v gelatin derived from bovine Type B (MilliporeSigma), was then stirred into the alginate for 2.5 hours at RT. For the development of the alginate/methylcellulose (Alg/MC) bioink, 3% w/v alginate was dissolved in DMEM/F12 for one hour at 35 °C. The alginate was then cooled to RT, mixing continuously for two hours. 6% w/v methylcellulose (MilliporeSigma) was added, mixing for 3 hours at 25 °C, and 0.5 hours at 50 °C to remove any air bubbles present. Ink was then transferred to 3 mL print cartridges and stored at 4 °C until use.

### Printing of Blank Alginate/Methylcellulose Bioink

Prior to printing, all bioinks were placed at 25 °C for 2 hours and UV sterilized for 1 hour. In the case of the alginate/gelatin bioink, it was placed at 37 °C for an additional five minutes, to prevent the printing of a granular hydrogel. The printer used to form all printed constructs was the BioX 3D extrusion bioprinter (Cellink). The ink was extruded through a 410 μm nozzle (Cellink), at pressures of 8-30 kPa, and nozzles speeds of 5-45 mm s^-1^, depending on the viscosity and yield pressure of the ink. The print cartridge maintained temperatures of 25-30 °C. The print construct was extruded onto either a glass (Cellink) or plastic Petri dish (Fisher Scientific), with all printed constructs embedded with cells printed onto a plastic Petri dish. Images were taken immediately after extrusion of the bioink with the Cellink BioX camera head. 50 or 100 × 10^-3^ M CaCl_2_ (MilliporeSigma) with 10 × 10^-3^ M HEPES (MilliporeSigma) was manually added to the surface of the printed construct for 5 minutes, to crosslink the printed structure. The construct was then imaged again using the BioX camera head. All images of the printed constructs pre- and post-crosslinking were analyzed in ImageJ, with the diameter of each construct quantified using pixel to distance scales.

### Oxygen and Nutrient Diffusion Characterization of Printed Construct

To test the oxygen concentration and diffusion within the printed construct over time, the Alg/MC bioink was manually mixed with oxygen-insensitive and oxygen-sensitive beads. 5 million DAPI loaded polystyrene (oxygen insensitive) beads (Fisher Scientific) and 5 million Ruthenium Tris(2,2′-bipyridyl) dichloride hexahydrate (RTDP) (oxygen sensitive) beads (Fisher Scientific) were both suspended in PBS (0.1 mL) (Gibco), and gently mixed into the alg/MC (0.5 mL) bioink.^[81]^ The ink was printed using a Cellink BioX with a 410 μm nozzle, at 25 °C, with a printing pressure of 25 kPa and a nozzle speed of 12 mm s^-1^, onto a plastic Petri dish. All printed constructs were immediately crosslinked with 50 × 10^-3^ M CaCl_2_ for 5 minutes. The printed structures were then incorporated into a micro physiological device in order to evaluate the oxygen concentration in printed structures.^[81]^

The device was then perfused with culture media loaded with 1 mg/mL of FITC labelled Dextran (70kDa) at a rate of 30 μL hr^-1^. The FITC signal from the perfused media was also obtained at regular intervals (30 min) over a period of 24 hours to probe the diffusion properties of the ink. 6 ROIs within the printed structures were analyzed for change in fluorescent intensity over time (Figure 2b). The fluorescence intensity ratio was calculated over time for each specific ROI by dividing the fluorescence intensity by the boundary fluorescence intensity.

The diffusion coefficient for 70 kDa Dextran in the printed constructs was calculated using the FITC fluorescence intensity ratio, and Fick’s Laws of Diffusion. Diffusion was assumed to be linear, and convective flow within the structure was considered to be negligible. It was also assumed that there was a constant concentration source and constant diffusion length. The distance between the ROI and the boundary of the structure was measured as the length that the dextran traveled. The location of all ROIs can be seen in Figure 2b. The generalized solution can be found below, where D is the diffusion coefficient (cm^2^ s^-1^), x (cm) is the distance between the boundary of the structure and the ROI, t (s) is time when the concentration gradient with respect to time is constant (steady state), I_ROI_ is the intensity at time t at the ROI, I_boundary_ is the constant intensity at the boundary of the structure, and erfc is the complimentary error function. The complementary error function was calculated using

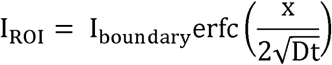

In parallel the oxygen concentration in printed structures was evaluated. ^[81]^ At the start of the experiment (t =0), the oxygen concentration is 18%, which is the ambient concentration of dissolved oxygen in the bioink and the perfused media. Oxygen concentration within the printed structure was monitored over the course of 24 hours under perfusion. The oxygen concentration in the printed structures was estimated using a fluorometric method by imaging the entire device at regular intervals for a period of 24 hours using Incell Analyser 6000 (GE Life Sciences) and measuring the intensities of the beads. This ratio of intensities is indirectly proportional to the amount of oxygen present in their environment as fluorescence signal by RTDP is quenched by oxygen.^[81]^ By determining the ratio of intensities at initial condition (18% oxygen) and at 0% oxygen (artificially created in the platform by introducing glucose oxidase), a standard curve for the intensity ratio and oxygen concentration is determined.

### Atomic Force Microscopy (AFM) Characterization of Printed Blank Alginate /Methylcellulose Bioink

Blank 3% w/v Alg/ 6% w/v MC stored at 4 °C was warmed for 2 hours at 25 °C before printing. The Alg/MC bioink was printed using a 410 μm nozzle (Cellink) at 25 kPa and 12 mm-s^-1^ onto a Superfrosted™ glass slide (Fisher Scientific). Further detailing for AFM construct setup and imaging can be found in Supplemental Materials and Methods. A force-volume map was recorded for each sample with a 25 x 25 μm^2^ area recorded, resulting in 256 measured indentations, with a maximum applied load of 2.75 nN. The force-indentation curves were processed using NanoScope Analysis software (Bruker) and the Young’s modulus from each of the force-indentation curve was determined using the Hertzian model for spherical indenter.^[82]^

### 3D Bioprinting and Culture of Single Cell-Embedded Bioink

Single cell printing was utilized to test the formation of aggregates within the printed construct for an extended culture period. iPSC-2’s were dissociated to single cell form (see Supplemental Material and Methods for further details). The ratio of cells to bioink volume used was 1.6 million cells per 1 mL of 3% w/v alg/ 6% w/v MC bioink. The single cells were added to a 1 mL syringe, which was attached to the print cartridge through a female-female LuerLock, and the ink and cells were gently mixed between the print cartridge and the syringe. The ink cartridge was transferred to the printer and attached to a 410 μm nozzle. Printing conditions were at 25 °C, with the printing pressure set at 25 kPa, and the nozzle speed set at 12 mm s^-1^. After printing the cell construct was crosslinked for 5 minutes using 50 × 10^-3^ M CaCl_2_ and incubated in mTeSR1 with 10 × 10^-6^ M Y-27632. Phase images were taken with an Olympus CKX41 at the same time each day over the 8-day print incubation period. The average aggregate diameter was quantified using ImageJ from Day 8 images.

### 3D Bioprinting and Culture of Aggregate-Embedded Bioink

Prior to printing, each type of cell was prepared for mixing within the bioink. Primary islets prior to printing were incubated in PIM(R) and for each printing experiment, approximately 500 aggregates were mixed in 3% w/v alg/ 6% w/v MC bioink (0.5 mL). All aggregates, primary or iPSC-derived, were printed in the same manner. Undifferentiated iPSC-1 and iPSC-2 aggregates were formed from aggregating single cells in in-house fabricated agarose microwells.^[40]^ For post-print differentiation experimentation, iPSC-2 aggregates at either the pancreatic progenitor, endocrine progenitor, or iPSC islet-like organoid stage were decapsulated using EDTA from the alginate bead previously described in the iPSC Culture and Encapsulation section.

All aggregates, primary or iPSC-derived, were added to the bioink, printed, and crosslinked in the same manner as the single cell bioprinting. Primary human islets were cultured in PIM(R), undifferentiated single cells and aggregates were cultured in mTeSR1 with 10 × 10^-6^ M Y-27632, and differentiating iPSC aggregates were cultured in the appropriate differentiation media (see Supplemental Information: iPSC Differentiation into Pancreatic Islet Endocrine Cells). For any printed constructs intended for GSIS, the structures were immediately moved to 12-well 3 μm pore size transwells (Corning).

As control for primary human islet studies, free-floating primary human islets were cultured in PIM(R) for the same duration as the printed primary islets. In the experiments involving printed iPSC-derived islets, control non-printed iPSC-derived islets were kept and maintained in the 1.5% w/v alginate beads for the same duration of culture as the printed iPSC-derived islets.

### MTS Characterization of Printed Cell Constructs

To quantify the viability of the cells embedded in the printed constructs, control and printed aggregates were collected for 4 hours and 3 days post-printing and tested following the CellTiter96 (Promega) assay protocol. Undifferentiated control iPSC-1 and iPSC-2 aggregates were taken from alginate beads (as discussed previously in iPSC Culture and Encapsulation section) and cultured for the same amount of time. Each sample (in triplicate) reading was normalized based on the number of aggregates in the sample. The printed constructs were further normalized by the absorbance reading of the control, so that the printed cell viability could be directly compared to the control cell.

### LDH Characterization of Printed Cell Constructs

To further quantify the undifferentiated cell viability, a CyQuant LDH cytotoxicity assay (Invitrogen) was conducted on both control aggregates and printed aggregates, following the protocol provided by the manufacturer. Both the control aggregates and the printed aggregates were cultured in mTeSR1with 10 × 10^-6^ M Y-27632. Every 24 hours during the desired cultured time of the printed construct, supernatant media (300 μL) was sampled from both the control alginate beads and the printed construct and stored at -80 °C until testing. The fluorescence intensity for each sample was measured using a SpectraMax M5 at an excitation of 560 nm and emission of 590 nm. Sample values were normalized using a blank control, and by the number of aggregates within each printed construct and the control alginate beads.

### Live/Dead Staining of Printed Cell Constructs

To visualize the printed cell viability, cells were tested with a Live/Dead fluorescence kit (Invitrogen). For printed constructs, cells were either decapsulated with EDTA prior to staining or left intact within the construct. All cells were stained and imaged for viability 1-8 days after printing. The live cell stain Calcein AM, and the dead cell stain Ethidium Homodimer-1 (Ethd-1) were diluted to 1 × 10^-6^ M in culture media. Both stains were added to the cells and incubated in the dark at room temperature. After 35 minutes, the excess stain was removed, and samples were imaged on an Olympus IX81 at 4x and 10x. Control primary human islets were stained and imaged following the same protocol as the printed primary human islets. Control iPSC-derived islets were decapsulated from the alginate beads with EDTA but then stained following the same protocol as the printed iPSC-derived islets. Aggregate size was quantified from 4x and 10x images using ImageJ.

Viability in islets and iPSC aggregates was determined by calculating the ratio of live cell area labelled green by Calcein AM to the sum of live and dead cell area using ImageJ. As Ethd-1 only labels the nucleus of dead cells, we used an empirically determined correction factor (ratio of cell area to nucleus area= 2.3) which was multiplied to the Ethd-1 labelled areas (dead cell nuclei) to estimate the actual dead cell area. The viability was then calculated as the total area labelled green (live cell area) divided by the sum of the area labelled green and the corrected red labelled area (dead cell area). The Ethd-1 positive regions (dead cell nuclei) in the islets/ iPSC aggregates were located using ImageJ and were plotted as a function of distance (r) with respect to the centroid of the islet/iPSC aggregate to determine the spatial distribution of dead cell population in all the islets/iPSC aggregates represented in terms of the islet/iPSC aggregate size (R). The ratio r/R was plotted to determine the regions where there are more dead cells. An r/R closer to 1 indicates dead cells at the edge/periphery of the islet/aggregate whereas r/R closer to 0 would indicate the presence of dead cells at the core of the islet/iPSC aggregate.

### Immunofluorescent Staining of Printed Cell Aggregates

After completing the desired culture time in the printed construct, the structures were dissolved using EDTA. The aggregates were then fixed overnight in sterile filtered 4% w/v Paraformaldehyde in PBS at 4 °C. All details regarding primary and secondary antibodies for all conditions can be found in Supplemental Material and Methods. Aggregates were mounted on a glass slide using Gold Antifade with DAPI before imaging on an Olympus Fluoview confocal microscope.

### PCR Analysis of Printed Cell Aggregate

RT-qPCR analysis was conducted so that the expression of key gene markers of the printed cells could be quantified. After three days within the printed construct, the aggregates were decapsulated from the construct using EDTA and fixed using RNAzol (500 μL) (Sigma Aldrich). The control aggregates followed the same protocol, with decapsulation from the 1.5% alginate beads. The RNEasy Plus Mini kit (Qiagen) protocol was used for mRNA isolation. cDNA was obtained using Improm II Reverse Transcription (Promega), with each PCR reaction containing 5 μL SYBR Green Master Mix (Agilent), 3 μL nuclease free water, 1μL primer, and 1 μL cDNA. Analysis followed the ΔΔC_t_ method for RT-qPCR, with GAPDH acting as the housekeeping gene. The gene expression of the printed aggregates was normalized based on the gene expression of the control aggregates, with three unique samples tested for the printed aggregates.

### Glucose Stimulated Insulin Secretion (GSIS) Analysis

To determine the primary islet and iPSC-2 islet-like organoid insulin response to glucose, GSIS was performed. For primary human islets, 3 printed constructs were used, and for iPSC-derived islets, 5 printed constructs were used. Each printed construct was a sample that was kept in individual 12-well 3 μm transwells (Corning) for 7 days of culture, in both the primary islet testing and the iPSC derived islet testing. Control primary human islets were kept in free-floating culture for 7 days, then transferred to 12-well 3 μm transwells. Control iPSC-derived islets were kept in 1.5% w/v alginate beads (described in iPSC Culture and Encapsulation) for 7 days, and then decapsulated using EDTA and placed in a 12-well 3 μm transwell. All printed and iPSC islet control samples were placed in low glucose conditions for 1.5 hours, where media was collected, and then these conditions were repeated, resulting in two distinct low glucose conditions. The samples were then placed in the high glucose G55 buffer for 1.5 hours, media collected, and this process repeated for two distinct high glucose conditions. Control primary islet samples were placed in low glucose G55 buffer for 1.5 hours, then moved to high glucose G55 buffer for 1.5 hours. Further details for GSIS media used and the protocol followed can be found in Supplemental Materials and Methods. All collected media was stored at 4 °C and used for insulin quantification with an Insulin ELISA (Alpco), conducted based on the manufacturers protocol. Insulin release values were normalized based on the number of islets or iPSC islet organoids within the printed construct.

### scRNASeq Processing and Data Analysis

After 7 days in culture, printed constructs were dissolved using EDTA, and iPSC islet aggregates were harvested and resuspended in DMEM/F12. In parallel, control iPSC islet aggregates cultured in alginate beads were also decapsulated. TrypLE was added to the for 15-20 minutes, and manual dissociation every five minutes broke the aggregates into single cell form. Samples were suspended in PBS + 10 × 10^-6^ M Y-27632 and sent to the Single Cell Core at the University of Pittsburgh for further quality control. Each sample was verified to meet the >70% viability threshold by using the Countess II Automatic Cell Counter (ThermoFisher). Library preparation and droplet sequencing was done according to the protocol set by 10X Genomics for a 3PrimeV3.1 experiment, using the Chromium NEXT Gem Single Cell 3’ Reagent Kit v3.1 (Dual Index). cDNA quality was assessed with the Agilent TapeStation High Sensitivity D5000 ScreenTape and reagents, while final libraries were assessed using Agilent TapeStation High Sensitivity D1000 ScreenTape and reagents.

After sequencing, count files were aligned to a human reference genome (refdata-gex-GRCh38-2020-A) using the 10X Genomics CellRanger pipeline (v7.1.0 including intronic reads). All data was then transferred to Partek Flow (Partek Incorporated) for further analysis. The samples were filtered for high mitochondrial expression and high read counts, to eliminate doublets and potential damaged cells from the analysis. The sample counts were then normalized and differential analysis using an ANOVA model was performed to generate a feature gene list with differential gene expression. The feature gene list (gene identifiers, log2 fold change, and p-values) was exported to Ingenuity Pathway Analysis (IPA; Ingenuity Systems, Qiagen) where the data set was further filtered to exclude any differentially expressed genes that did not meet the threshold of p <0.05, and an absolute value Log2Ratio <1. Canonical pathway analysis was used to identify any biologically relevant pathways that were significant between the printed sample and the control sample. To determine the effects of printing stress specifically on the endocrine cell population, only cells that had expression levels of *INS*, *GCG*, and *SST* > 0 were filtered from the general cell population. An ANOVA was used to analyze differential gene expression between the filtered control and printed iPSC endocrine cell population for specific stress and mechanosignaling genes, and only genes with p < 0.05 were considered to be differentially expressed. Violin plots were then generated that quantitatively compared gene expression between the control group and printed islet data set.

### Statistical Analysis

All quantitative data was expressed as mean ± standard deviation, unless stated otherwise, and plotted using GraphPad Prism 9. Data were tested for statistical differences using multiple t-tests, where *: p < 0.05, **: p < 0.01, ***: p < 0.005, and ****: p < 0.001 were considered statistically significant.

## Acknowledgements

M.P. has been supported by the U.S. Department of Education, Graduate Assistance in Areas of National Need (GAANN) Program, award number P200A180097. This work was supported by the following grant from the National Science Foundation (NSF): 2229156-NSF/CBET. Prashant N. Kumta would also like to acknowledge the Edward R. Weidlein Endowed Chair Professorship funds for the support of this work. We thank Dr. Alejandro Soto-Gutierrez from the University of Pittsburgh for generating and providing the hfl262 cell line. We thank Dr. Daniel Lamont for assistance with AFM. All AFM experimentation was conducted in The Nanoscale Fabrication and Characterization Facility (NFCF) in the Gertrude E. & John M. Petersen Institute of NanoScience and Engineering (PINSE) at the University of Pittsburgh. We thank Heidi Monroe and the University of Pittsburgh Single Cell Core Facility (RRID: SCR_025110) for the Illumina RNAsequencing. We thank Sri Chaparala and Dr. Ansuman Chattopadhyay for their help in the bioinformatics analysis. Partek Flow and Ingenuity Pathway Analysis software licensed through the Molecular Biology Information Service of the Health Sciences Library System, University of Pittsburgh was used for data analysis.

M.P. conceptualized and designed the experiments under the supervision of I.B. and P.N.K.. R.M.F., Z.L., and A.S.G. conducted the hfl262 cell characterization experiments. B.M., R.W. participated in the bioprinting experiments. C.W. participated in cell culture and GSIS. R.K. participated in the immunofluorescent microscopy and nutrient diffusion studies. M.P. wrote the manuscript under the supervision of I.B. and P.N.K..

