## Supplemental Information for "Bioprinting of Human Primary and iPSC-derived Islets with Retained and Comparable Functionality"

**Supplemental Material and Methods**

*hFL262 iPSC Generation and Characterization*

Primary human fetal fibroblasts were isolated as previously described.^[1]^ Briefly, the tissue was digested in EMEM (Lonza), which contains 0.5 mg ml^-1^ of collagenase Type XI (SigmaAldrich), on a lab shaker for 40 minutes. Viability was assessed by Τrypan blue exclusion test and was routinely >85%. Fetal fibroblasts were plated at a density of 1.3x10^5^ cells cm^-2^ on type I rat tail collagen coated 12 well plates (Corning). Cells were cultured and passaged 2 times to get a 100% pure population of human fetal fibroblasts, with a DMEM medium (Gibco) containing 1x Pen/Strep, × 10^-7^ m of insulin (Sigma-Aldrich), and 5% bovine serum albumin (Gibco). Reprograming of fetal fibroblasts was performed using episomal plasmids vectors adapted from a previously described protocol.^[1]^ Briefly, for each nucleofection, 1 million cells were resuspended in 100 μL of the AmaxaTM NHDF Nucleofector kit (Lonza), containing 3 μg of each of the four episomal plasmids vectors encoding OCT3/4 and p53 shRNA, SOX2 and KLF4, L-MYC and LIN28, and enhanced green fluorescent protein (eGFP) (Addgene). Cells were nucleofected using the Amaxa 4D- Nucleofector (Lonza) and plated in mTeSR1^TM^ on human embryonic stem cell–qualified Matrigel (Corning)-coated plates. hFL262 iPSC was expanded and pluripotency validated by mRNA expression of *OCT3/4*, *NANOG*, *LIN28A*, *SOX2* and *cMyc* using iPSCs derived from human Amniotic Epithelial cell (hAE iPSC) as positive control (Supplemental Figure 8a).^[2]^ Protein expression of the stemness marker OCT3/4, NANOG, SSEA4 and TRA-1-60 was assessed by immunofluorescence (Supplemental Figure S8b).

*iPSC Culture and Encapsulation*

To obtain a single cell suspension, undifferentiated iPSCs were incubated in mTeSR1 (STEMCELL Technologies) with 10 × 10^-6^ m Y-27632 (R&D Systems) overnight prior to encapsulating. To dissociate, the cells were incubated with Accutase (STEMCELL Technologies) for 7 min at 37 °C to detach cells and pipetted up and down obtain a single cell suspension. Cells were suspended in 1.5% (w/v) low viscosity alginate (MilliporeSigma) at room temperature at a ratio of 1 million cells: 3 mL alginate and added dropwise to a bath of 100 × 10^-3^ m calcium chloride (MilliporeSigma) with 10 × 10^-3^ m HEPES (MilliporeSigma) using a 22-gauge needle. Alginate capsules were incubated for 5 min in the 100 × 10^-3^ m calcium chloride and 10 × 10^-3^ m HEPES solution at room temperature to allow for complete crosslinking and gelation of the alginate capsules. Capsules were washed three times with PBS and suspended in mTeSR1 with 10 × 10^-6^ m Y-27632. Before starting differentiation, encapsulated cells were cultured for 4 days in mTeSR1 with 10 × 10^-6^ m Y-27632 followed by 1 day of mTeSR1.

*iPSC Differentiation into Pancreatic Islet Endocrine Cells*

The differentiation base media is composed of 2.44 × 10^-3^ m D-Glucose (Gibco), 1.23 g L^-1^ NaHCO_3_ (MilliporeSigma), 2% FAF-BSA (Fisher Scientific), 2 × 10^-3^ m Glutamax (Gibco), 1% Pen/Strep (Lonza), and MCDB131 (490 mL) (Corning/Gibco). Media changes were completed as follows with supplements to the differentiation base media. Definitive Endoderm Media (Days 1-3): differentiation base media was supplemented to have 2.46 g L^-1^ NaHCO_3_, 0.25 × 10^-3^ m Vitamin C (MilliporeSigma), and 1:50 ITS-X (MilliporeSigma). Day 1: 100 ng mL^-1^ Activin A (R&D Systems) and 1.4 μg mL^-1^ Chir99021 (Stemgent). Days 2-3: 100 ng mL^-1^ Activin A. Primitive Gut Tube (Days 4 and 6): 50 ng mL^-1^ KGF (Peprotech), 0.25 × 10^-3^ m Vitamin C, and 1:50 ITS-X. Pancreatic Progenitor 1 (Days 7 and 8): 50 ng mL^-1^ KGF, 0.25 × 10^-6^ m Sant1 (MilliporeSigma), 2 × 10^-6^ m Retinoic Acid (MilliporeSigma), 500 × 10^-9^ m PdBU (MilliporeSigma), 0.25 × 10^-3^ m Vitamin C, 10 × 10^-6^ m Y-27632, and 1:200 ITS-X. Pancreatic Progenitor 2 (Days 9, 11, and 13): 50 ng mL^-1^ KGF, 0.25 × 10^-6^ m Sant1, 0.1 × 10^-6^ m Retinoic Acid, 0.25 × 10^-3^ m Vitamin C, 10 × 10^-6^ m Y-27632, 5 ng mL^-1^ Activin A and 1:200 ITS-X. Endocrine Progenitor (Days 14-30): increase NaHCO_3_ to 1.75 g mL^-1^ and increase glucose to 20 × 10^-3^ m. Days 14 and 16: 0.25 × 10^-6^ m Sant1, 0.1 × 10^-6^ m Retinoic Acid, 0.25 × 10^-3^ m Vitamin C, 1 × 10^-6^ m XXI (MilliporeSigma), 10 × 10^-6^ m ALk5i II (Axxora), 1 × 10^-6^ m T3 (MilliporeSigma), 20 ng mL^-1^ Betacellulin (Fisher Scientific), 10 μg mL^-1^ Heparin (MilliporeSigma), 1:200 ITS-X. Days 18-30: 0.025 × 10^-6^ m Retinoic Acid, 0.25 × 10^-3^ m Vitamin C, 1 × 10^-6^ m XXI, 10 × 10^-6^ m ALk5i II, 1 × 10^-6^ m T3, 20 ng mL^-1^ Betacellulin, 10 μg mL^-1^ Heparin, 1: 200 ITS-X. Maturation (Days 32+): 1: 200 ITS, and 0.25 × 10^-3^ m Vitamin C.

*Atomic Force Microscopy (AFM) Characterization of Printed Blank Alginate/Methylcellulose Bioink*

UV adhesive (Norland Optical Adhesives, NOA72) was applied to glass slide using a 22G needle as a hydrophobic barrier around the printed construct, to help form the meniscus needed for AFM working in contact mode in liquid phase. PBS was also applied on the surface of the printed construct to prevent drying. A colloidal spherical borosilicate tip with a radius 5 μm (CP-qp-CONT-BSG, NanoAndMore USA) attached to a cantilever was used to probe the printed structures. All measurements were conducted using a Bruker Dimension Icon AFM (Bruker), which was controlled by a Nanoscope VI (Bruker). The spring constant of the cantilever was measured using the thermal fluctuation method.^[3]^ The measured spring constant of the cantilever was 0.13 N m^-1^. The deflection sensitivity for the colloidal tip was calibrated to be 41.65 nm V^-1^, using a standard sapphire substrate.

*Immunofluorescent Staining of Printed Cell Aggregates*

Primary antibodies for OCT4 (Mouse, R&D Systems) and NANOG (Goat, R&D Systems) along with secondary antibodies anti-mouse Alexa Fluor 647 and anti-goat Alexa Fluor 488 were used for undifferentiated iPSC aggregate imaging. Primary antibodies for NKX6.1 (Mouse, R&D Systems), PDX1 (Goat, R&D Systems), C-peptide (Mouse, R&D Systems), and Glucagon (Rabbit, Abcam) along with secondary antibodies anti-mouse Alexa Fluor 647, anti-goat Alexa Fluor 488, and anti-rabbit Alexa Fluor 555 were used for primary and iPSC-derived islet imaging. Primary antibodies were added at 1:100 and secondary antibodies were added at 1:500.

*Glucose Stimulated Insulin Secretion (GSIS) Analysis*

All control and printed samples were exposed to two periods of low glucose (3.0 × 10^-3^ m) or high glucose (16 × 10^-3^ m). The GSIS base culture media is low glucose G55 buffer, which is composed of DMEM-no glucose (500 mL) (Gibco), Hams F10 (Cytvia/Gibco), sodium bicarbonate (60 mg) (MilliporeSigma), calcium chloride (110 mg) (MilliporeSigma), and 2.5% FAF-BSA (Fisher Scientific). Before incubation, samples were washed 5 times in low glucose G55 buffer. For the primary human islets, all samples were incubated overnight in the low glucose G55 buffer. For iPSC-derived islets, all samples were incubated for 2 hours in low glucose G55 buffer. After the pre-incubation samples were washed 5x in low glucose G55 buffer.

**Supplemental Figures**

**Table S1.** Comparative functionality studies for islet extrusion bioprinting.

| **Cell Type** | **Printing Modality** | **Bioink Composition** | **Duration of Culture before GSIS** | **Results** | **Reference** |
| --- | --- | --- | --- | --- | --- |
| Primary Human Islets | Indirect Extrusion | 4 mg/mL type 1 rat tail collagen + 100 mg/mL fibronectin + 100 mg/mL collagen IV | 10 days, islets decapsulated | - Primary human islets in ECM gel were embedded into bioprinted PLGA scaffolds - After 10 days islets decapsulated and exhibited SI = 1.8 | Daoud et. al. (2011)^[4]^ |
| Primary Human Islets | Direct Extrusion | 4% w/v alginate/ 5% w/v gelatin | 7 days, islet intact within print | - Primary islets remained viable and functional after 1 day in culture, but lost functional after 7 days  -Functionality was restored post decapsulation from the print | Marchioli et. al. (2015)^[5]^ |
| Mouse Islets | Direct Extrusion + Photoinitiated Crosslinking | 2% w/v alginate + 7.5% w/v GelMA | 3 days, islet intact within print | - The printed mouse islets did not respond appropriately to glucose, resulting in an SI < 2 | Liu et. al. (2019)^[6]^ |
| Primary Human Islets + Human Insulin Producing iPSCs | Indirect Extrusion | 2% w/v pancreatic decellularized ECM | 5 days, islet intact within scaffold | -Primary human islets incorporated into the bioink maintained an SI of 3.1 after 5 days (not printed)  -Insulin producing iPSCs incorporated into the bioink, and cultured for 10 days demonstrated a constant insulin level | Kim et. al. (2019)^[7]^ |
| Human Embryonic Stem Cells | Direct Extrusion | 1% w/v pancreatic decellularized ECM printed into PCL capsule | Not reported | -GSIS indicated that printed aggregate ESC derived insulin producing cells responded appropriately to varying glucose levels | Hwang et. al. (2021)^[8]^ |
| Rat Islets | Direct Extrusion | 3% w/v alginate/ 9% w/v methylcellulose | 7 days, islet intact within print | - Printed rat islets after 7 days of culture remained viable and positively expressed insulin and glucagon  -GSIS indicated that the islets were the most functional (SI= 3.4) on Day 4 post print, but SI was reduced to 1.8 after 7 days culture | Duin et. al. (2019)^[9]^ |
| Neonatal porcine islet like cell clusters (NICCs) | Direct Extrusion | 3% w/v alginate/ 9% w/v methylcellulose | 21 days, islet intact within print | -NICC need several weeks to mature and display functionality  - NICCs supplemented with PBS, 1% BSA, plate lysate, or fresh frozen plasma all had an SI above 2 by 21 days post printing | Duin et. al. (2022)^[10]^ |
| Primary Human Islets + Adipose Derived Stem Cells | Direct Extrusion | 20% w/v alginate nano-fibrillated cellulose bioink | 14 days, islet intact within scaffold | -Human islets alone were contrasted to human islets with ASCs (1.2 x 10^6^/scaffold)  - Islets alone by Day 8 had reduced SI (0.7203) to islets + ASCs SI (2.4) | Abadpour et. al. (2023)^[11]^ |
| Mouse embryonic pancreatic progenitor cells | Direct Extrusion | Fibrin + 250 mg/mL gelatin type A + 3 mg/mL hyaluronic acid + 100 KIU aprotinin | GSIS not conducted but pancreatic progenitor ESCs cultured 7 days | -Mouse ESCs could successfully differentiate within the construct to endocrine, acinar, and ductal lineages, but did not fully mature | Edri et. al. (2024)^[12]^ |
| Human Embryonic Stem Cells | Direct Extrusion | 10 mg/mL pdECM + 0.1 mg/mL laminin + 0.1 mg/mL type IV collagen | 4 days, islet intact within scaffold | -ESC derived islets were printed and after 4 days of culture responded appropriately to glucose, resulting in SI ~ 3 | Kim et. al. (2025)^[13]^ |

**Table S2a:** Alginate at varying weight per volume concentrations was mixed with DMEM/F12. The bioink was then printed at different pressures and nozzle speeds and deposited on either a glass or plastic substrate. The percent difference in diameter between the plastic and glass substrate at the same speed and nozzle pressure was quantified by subtracting the diameter of the glass print from the plastic print and dividing by the diameter of the plastic print.

| Alginate Concentration (% w/v) | Pressure (kPa) | Speed (mm s^-1^) | Average diameter of plastic structure in comparison to glass structure |
| --- | --- | --- | --- |
| 3 | 8 | 20 | -35% |
| 3 | 8 | 30 | -29% |
| 3 | 5 | 40 | -44% |
| 4 | 15 | 20 | -86% |
| 4 | 8 | 20 | -7% |
| 4 | 8 | 30 | -21% |
| 4.5 | 15 | 20 | -43% |
| 4.5 | 8 | 20 | -47% |
| 4.5 | 8 | 30 | -62% |
| 5 | 15 | 20 | -48% |
| 5 | 8 | 20 | -49% |
| 5 | 8 | 30 | -48% |

**Table S2b:** Alginate at varying weight per volume concentrations was mixed with DMEM/F12. The bioink was then printed at different pressures and nozzle speeds and deposited on either a glass or plastic substrate. The percent difference in diameter between each alginate concentration print was determined following the same method utilized in Tables S1a and S1c.

| Pressure (kPa) | Speed (mm s^-1^) | Surface | Diameter of 4% w/v Alg print in comparison to 3% w/v alg | Diameter of 5% w/v Alg print in comparison to 3% w/v alg | Diameter of 5% w/v Alg print in comparison to 4% w/v alg |
| --- | --- | --- | --- | --- | --- |
| 15 | 20 | glass | -26% | -24% | 2% |
| 8 | 20 | glass | -30% | -20% | 15% |
| 8 | 30 | glass | -22% | -16% | 8% |
| 5 | 40 | glass | -24% | -19% | 6% |
| 15 | 20 | plastic | -35% | -47% | -17% |
| 8 | 20 | plastic | -34% | -37% | -5% |
| 8 | 30 | plastic | -21% | -38% | -22% |
| 5 | 40 | plastic | -13% | -49% | -55% |

**Table S2c:** Alginate at 4% w/v was mixed with either DMEM/F12 or DI water and printed at the same range of pressures and nozzle speeds, and then deposited on a glass or plastic substrate. The percent difference in diameter between the ink mixed with DMEM/F12 or DI water at the same speed and nozzle pressure was quantified by subtracting the diameter of the DMEM/F12 print from the DI water print and dividing by the diameter of the DI print.

| Concentration of Alg % | Pressure (kPa) | Speed (mm s^-1^) | Surface | Diameter of DMEM/F12 print in comparison to DI print |
| --- | --- | --- | --- | --- |
| 4 | 15 | 20 | glass | -22% |
| 4 | 8 | 20 | glass | -29% |
| 4 | 8 | 30 | glass | -26% |
| 4 | 5 | 40 | glass | -29% |
| 4 | 15 | 20 | plastic | -1% |
| 4 | 8 | 20 | plastic | -5% |
| 4 | 8 | 30 | plastic | 1% |
| 4 | 5 | 40 | plastic | 5% |

**Figure S1: Window of printability and strand formation for 4% w/v alginate, mixed in DMEM/F12, and printed at a range of pressures and nozzle speeds on a glass substrate.**

Pressure (kPa)

| Speed (mm/s) | Strand |
| --- | --- |
| 20 | 6 |
| 25 | 6 |
| 30 | 6 |
| 35 | 7 |
| 40 | 8 |
| 45 | 8 |
| 50 | 10 |

| Speed (mm/s) | Uniform Droplet |
| --- | --- |
| 25 | 5 |
| 30 | 5 |
| 35 | 5 |
| 40 | 7 |
| 50 | 8 |

| Speed (mm/s) | Non Standard Droplet |
| --- | --- |
| 50 | 8 |
| 20 | 5 |
| 40 | 7 |

**Figure S1:** Window of printability and strand formation for 4% w/v alginate, mixed in DMEM/F12, and printed at a range of pressures and nozzle speeds on a glass substrate.

**Table S2d:** 4% w/v alginate, 4% w/v alginate/ 4% w/v gelatin, and 3% w/v alginate/ 6% methylcellulose (MC) was printed at various pressures and nozzles speeds on either glass or plastic substrates. The diameter of the construct was then collected before and after crosslinking with either 50 × 10^-3^ m CaCl_2_ or 100 × 10^-3^ m CaCl_2_. The percent difference in diameter before and after crosslinking was determined following the same method utilized in Tables S1a, S1b, and S1c.

| Alginate concentration (w/v %) | Ink addition (w/v%) | Substrate | Pressure (kPa) | Speed (mm s^-1^) | Crosslinking concentration (× 10^-3^ m CaCl_2_) | Time Crosslinked | Diameter change after crosslinking |
| --- | --- | --- | --- | --- | --- | --- | --- |
| 4 |  | glass | 8 | 20 | 50 | 5 min | -25% |
| 4 |  | glass | 8 | 20 | 100 | 5 min | -26% |
| 4 |  | plastic | 8 | 20 | 50 | 5 min | -6% |
| 4 |  | plastic | 8 | 20 | 100 | 5 min | -23% |
| 4 | 4% w/v gel | glass | 30 | 8 | 50 | 5 min | 41% |
| 4 | 4% w/v gel | glass | 30 | 8 | 100 | 5 min | 26% |
| 4 | 4% w/v gel | plastic | 30 | 8 | 50 | 5 min | 5% |
| 4 | 4% w/v gel | plastic | 30 | 8 | 100 | 5 min | 19% |
| 3 | 6% w/v MC | plastic | 30 | 12 | 50 | 3 min | -4% |
| 3 | 6% w/v MC | plastic | 30 | 12 | 100 | 3 min | -6% |
| 3 | 6% w/v MC | plastic | 30 | 12 | 50 | 5 min | -12% |
| 3 | 6% w/v MC | plastic | 30 | 12 | 100 | 5 min | -6% |
| 3 | 6% w/v MC | plastic | 30 | 12 | 50 | 10 min | -7% |
| 3 | 6% w/v MC | plastic | 30 | 12 | 100 | 10 min | -5% |
| 3 | 6% w/v MC | glass | 30 | 12 | 50 | 3 min | -27% |
| 3 | 6% w/v MC | glass | 30 | 12 | 50 | 5 min | -22% |
| 3 | 6% w/v MC | glass | 30 | 12 | 50 | 10 min | -9% |

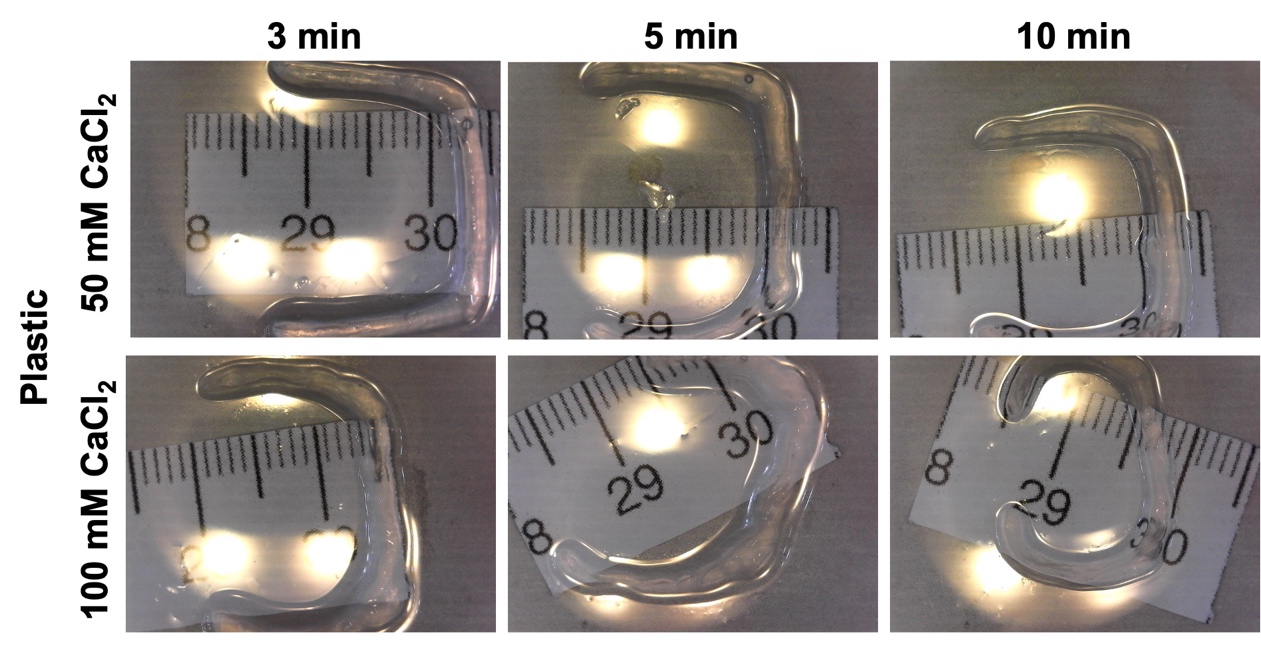

**Figure S2:** 3% w/v alginate/ 6% methylcellulose (MC) was printed at 30 kPa and 12 mm s^-1^ on a plastic substrate. The printed constructs were crosslinked with either 50 × 10^-3^ m CaCl_2_ or 100 × 10^-3^ m CaCl_2_ for 3, 5, or 10 minutes.

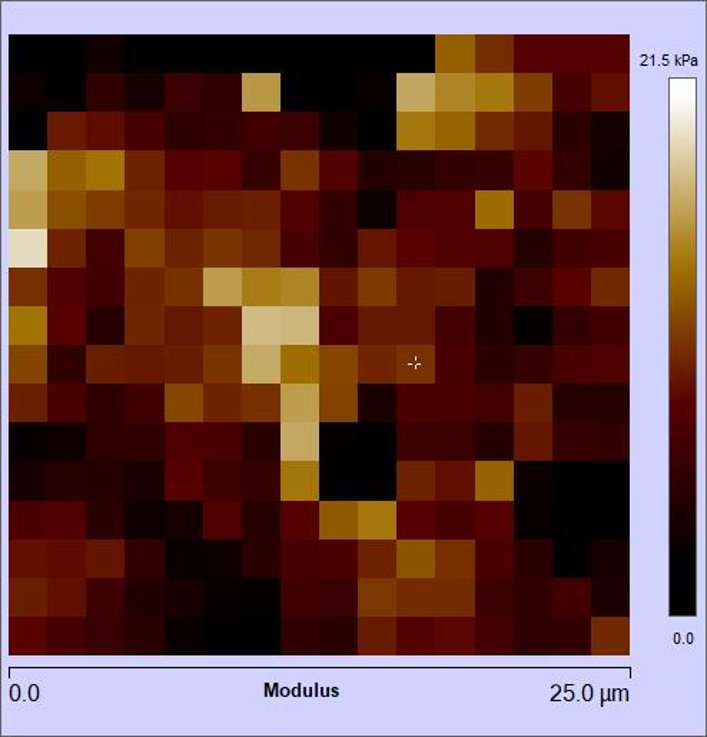

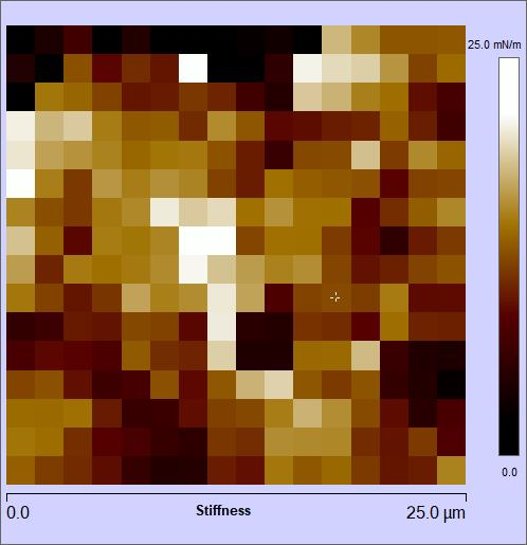

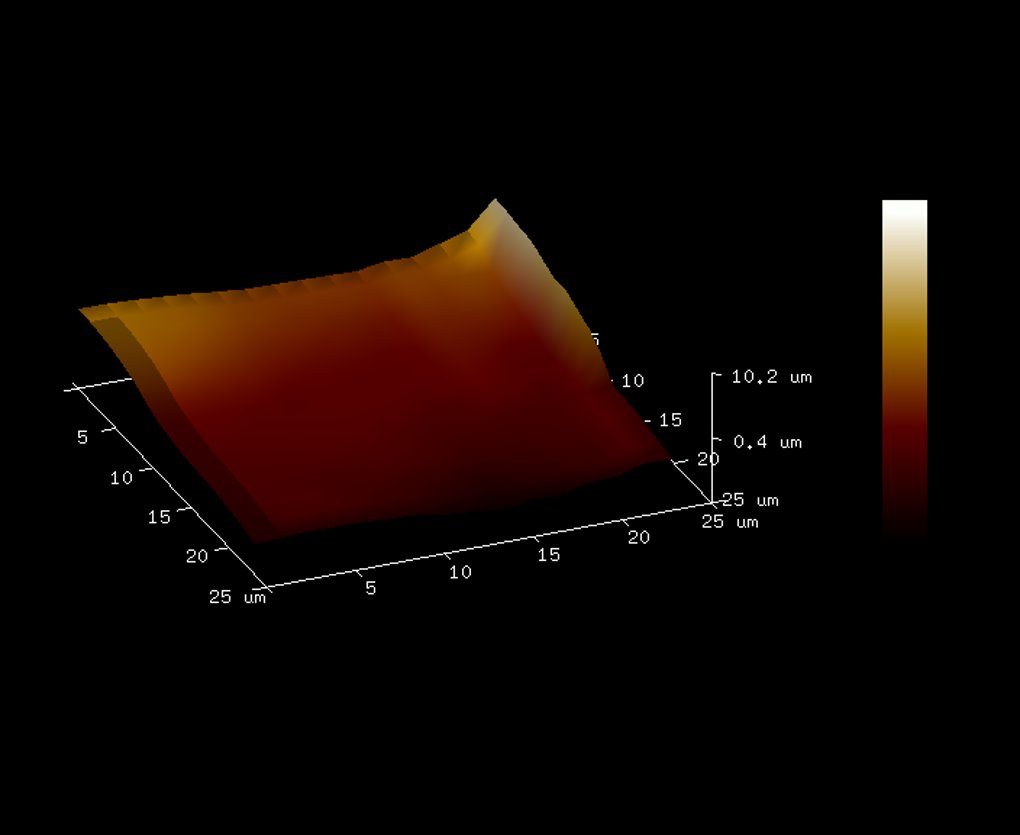

**A**

**B**

**C**

**Figure S3. A)** Representative **f**orce-volume elastic moduli heat map (25 x 25 μm) of printed construct measured using AFM contact mode in liquid. Printed construct was crosslinked with 50 mM CaCl_2_ and kept hydrated with PBS. **B)** Representative force-volume stiffness heat map (25 x 25 μm) of printed construct measured using AFM contact mode in liquid. **C)** Basic heat map of surface topography of printed construct. Heat map is 25 x 25 μm, and construct was hydrated with PBS.

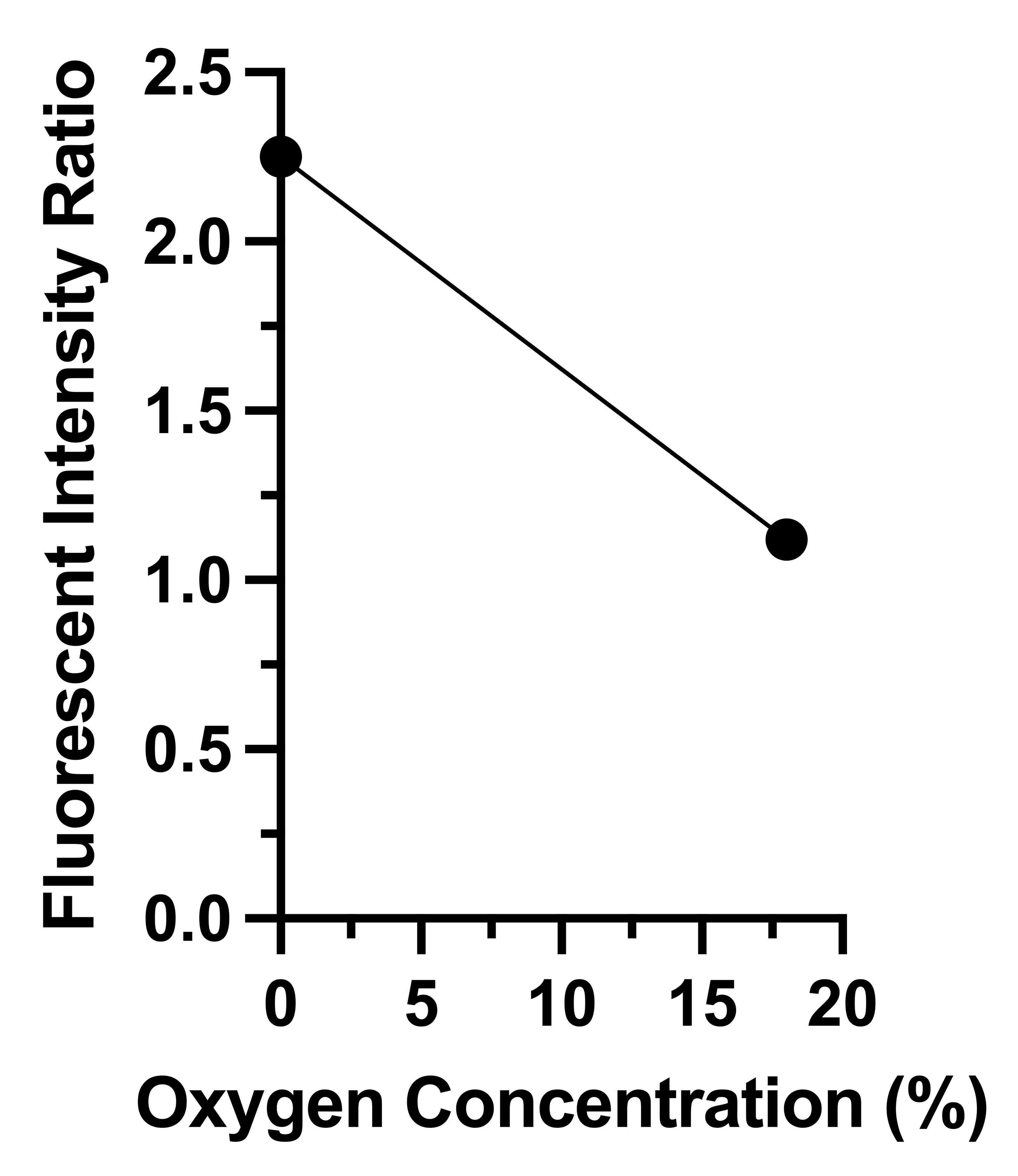

**Figure S4:** Linear calibration curve for converting fluorescent intensity ratio of RTDP (oxygen sensitive) and DAPI (oxygen insensitive) beads to oxygen concentration (%). As the fluorescent intensity ratio decreases, oxygen concentration increases.

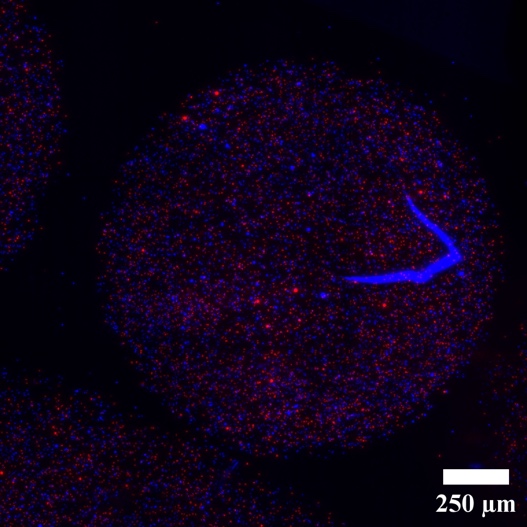

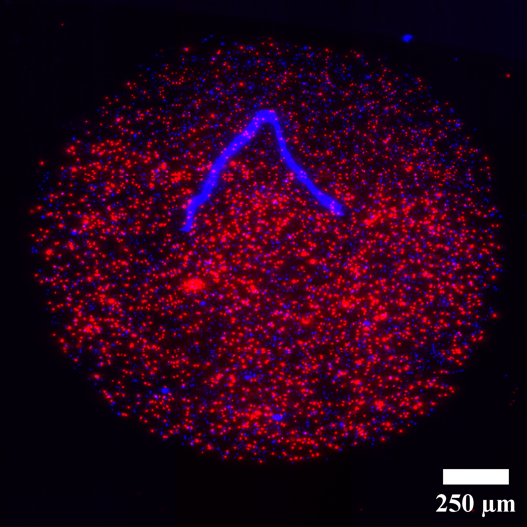

**Figure S5:** On left: 0% oxygen and corresponding fluorescence in RTDP beads in representative imaging condition. On right: 18% oxygen and corresponding fluorescence in RTDP beads in representative imaging condition.

**Table S3:** Donor statistics for primary cadaveric human islets used in experiments.

| **Category** | **Donor Statistics** |
| --- | --- |
| Sex | Male |
| Race | African American |
| Height | 70” |
| Weight | 206 lbs |
| BMI | 30.0 |
| A1c | 5.6 |

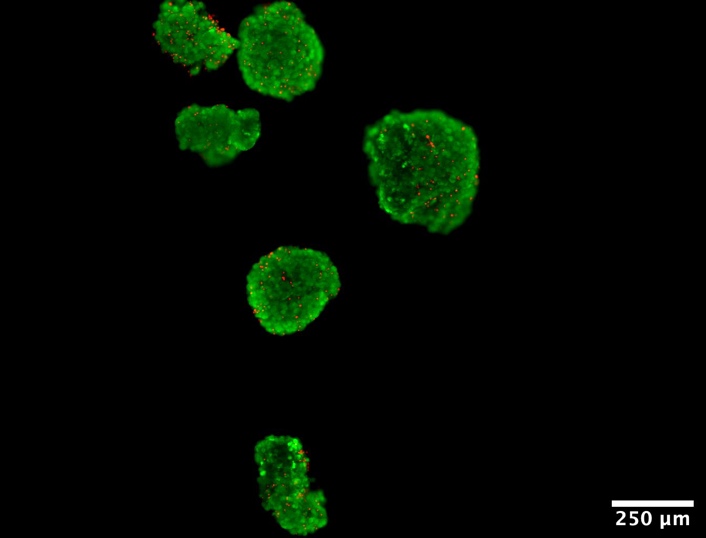

**Figure S6.** Representative LiveDead imaging of free-floating control primary human islets. Live cells are shown in green, dead cells are shown in red. Scale bar represents 250 μm.

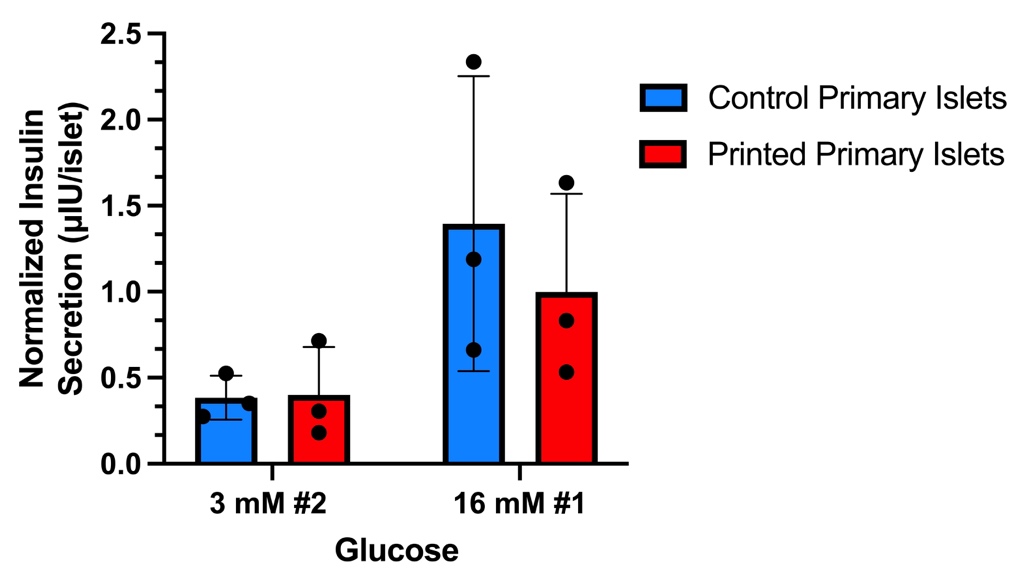

**Figure S7.** Glucose stimulated insulin secretion of printed and control primary human islets after 7 days in culture. Results for control primary islets are shown in blue, and results for printed primary human islets are shown in red.

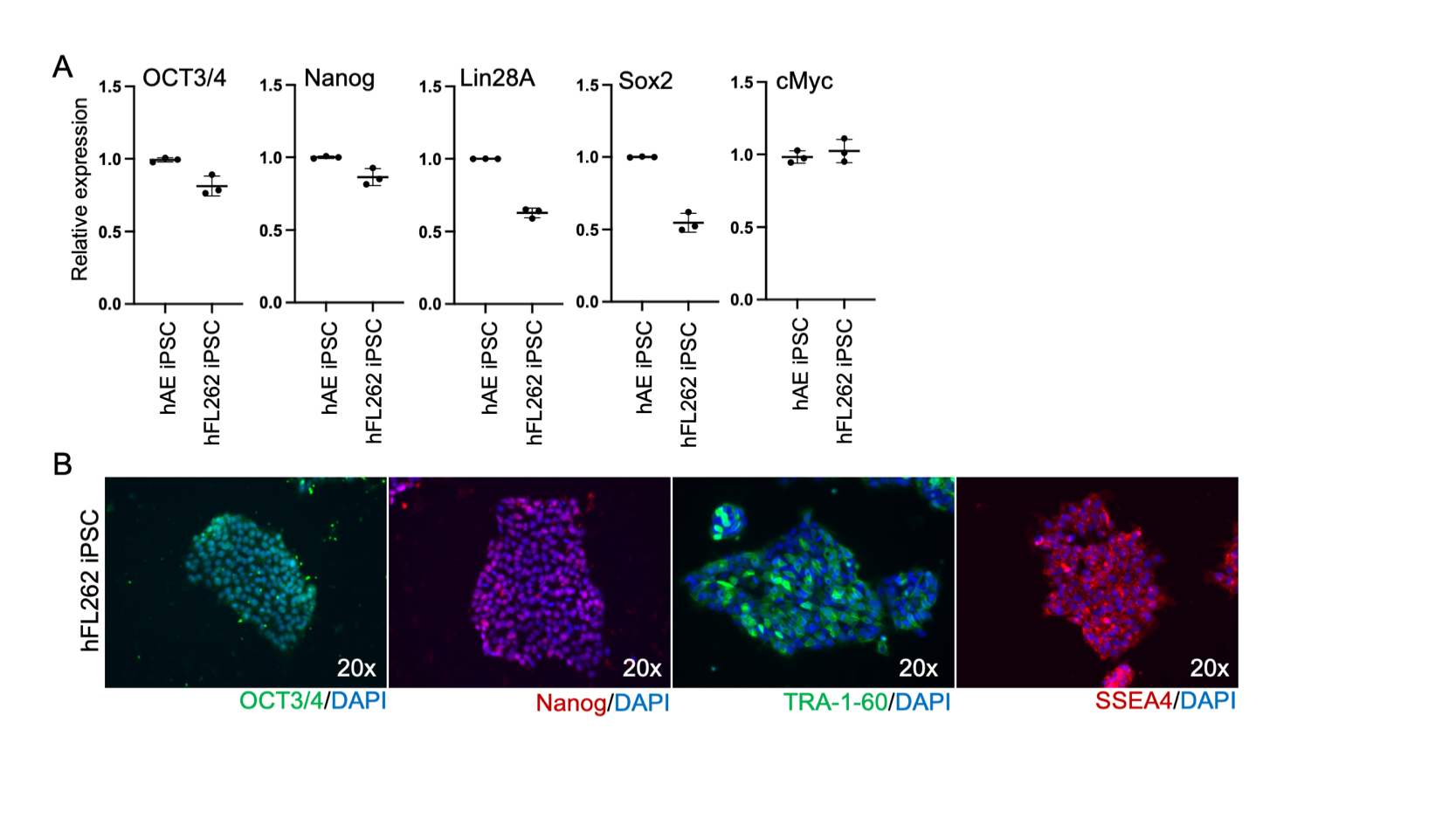

**Figure S8:** A) Quantitative gene expressions of pluripotency markers: *OCT3/4, NANOG, Lin28A, SOX2 and cMyc* in the hFL262 iPSC (n=3). iPSC derived from human Amniotic Epithelial cell (hAE iPSC) was used as a positive control (n=3). Values were determined relative to β-actin and presented as fold change relative to the expression in human hAE iPSC, which is set as 1. B) Immunofluorescence micrographs of pluripotency markers: OCT3/4, NANOG, SSEA4 and TRA-1-60 in the hFL262 iPSC.

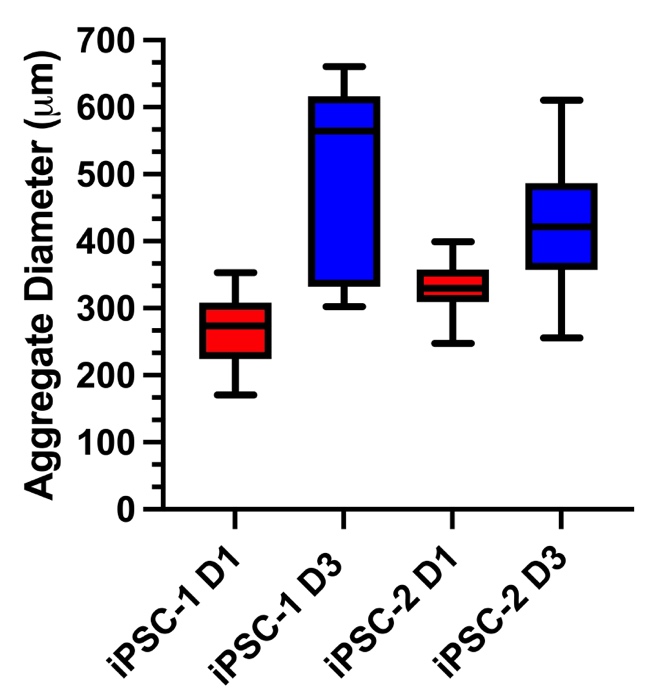

**Figure S9:** Aggregate diameter size distribution for undifferentiated iPSC-1 and -2 aggregates 1 and 3 days after printing.

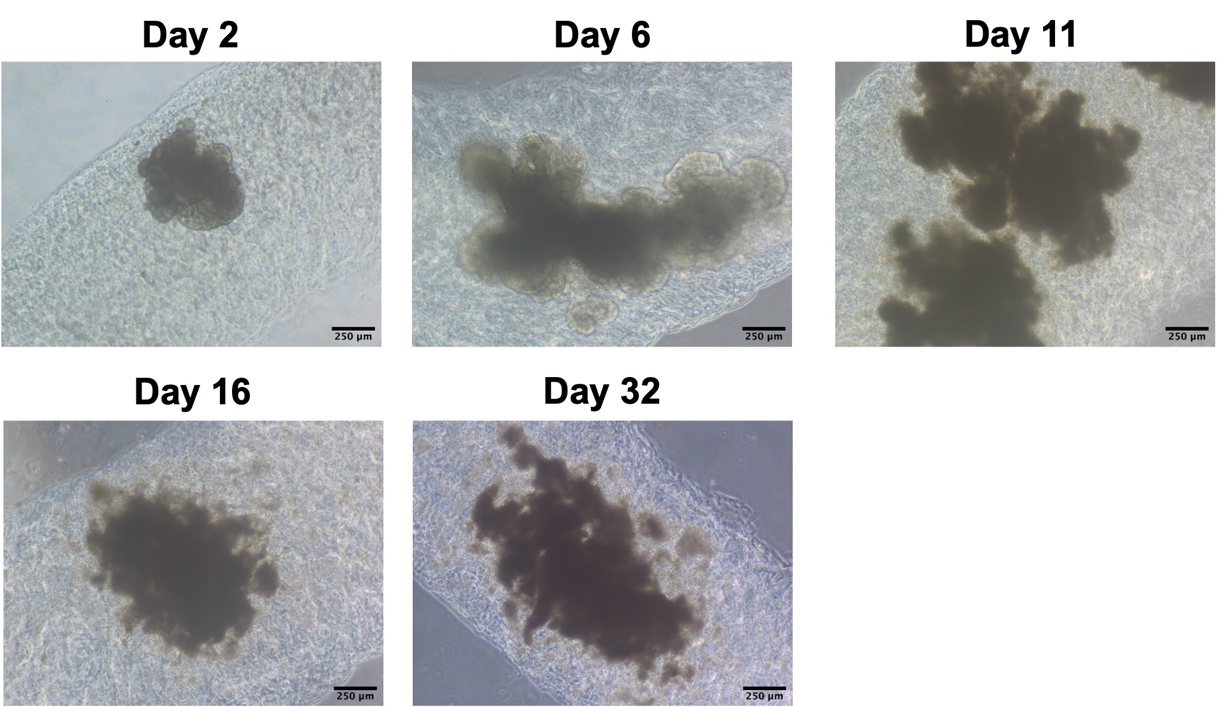

**Figure S10:** Undifferentiated iPSC-2 aggregates were printed and differentiated over 32 days to the immature iPSC islet stage. Phase imaging was taken on days 2, 6, 11, 16, and 32 to track aggregate growth and morphology. Scale bar representing 250 μm.

**Post-Print PP**

**Day 1**

**Day 14**

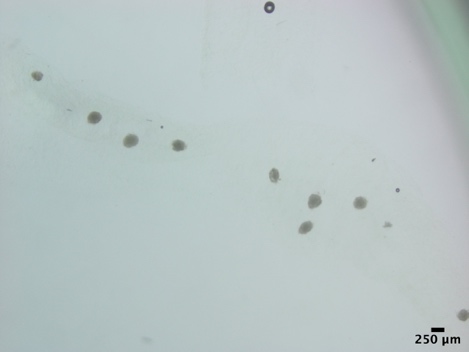

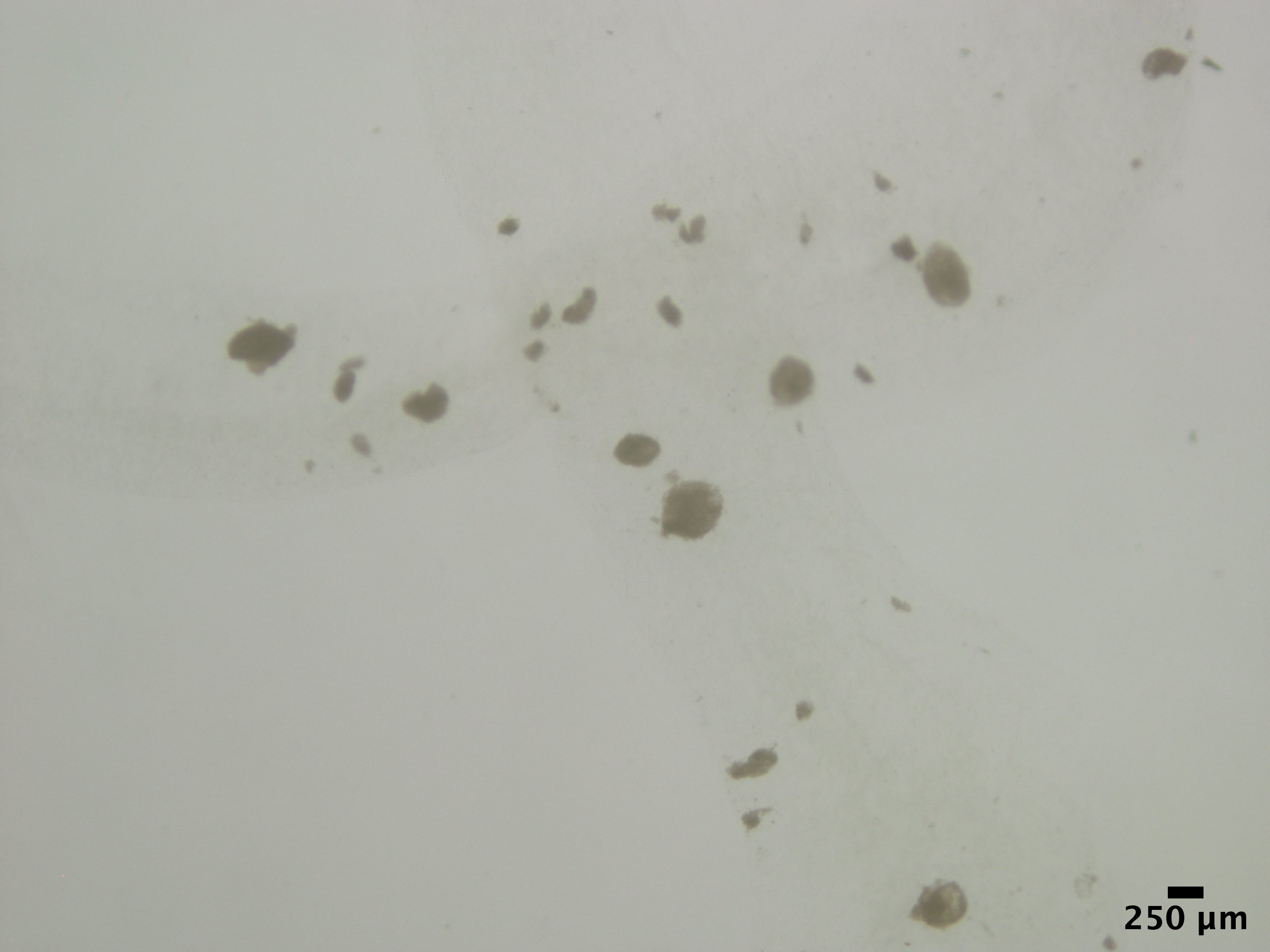

**Figure S11:** For post-print PP, undifferentiated iPSC aggregates were printed and then cultured for 14 days to the pancreatic progenitor stage. Phase imaging for post-print PP was taken on Days 1 and 14 post-printing. Scale bar representing 250 μm.

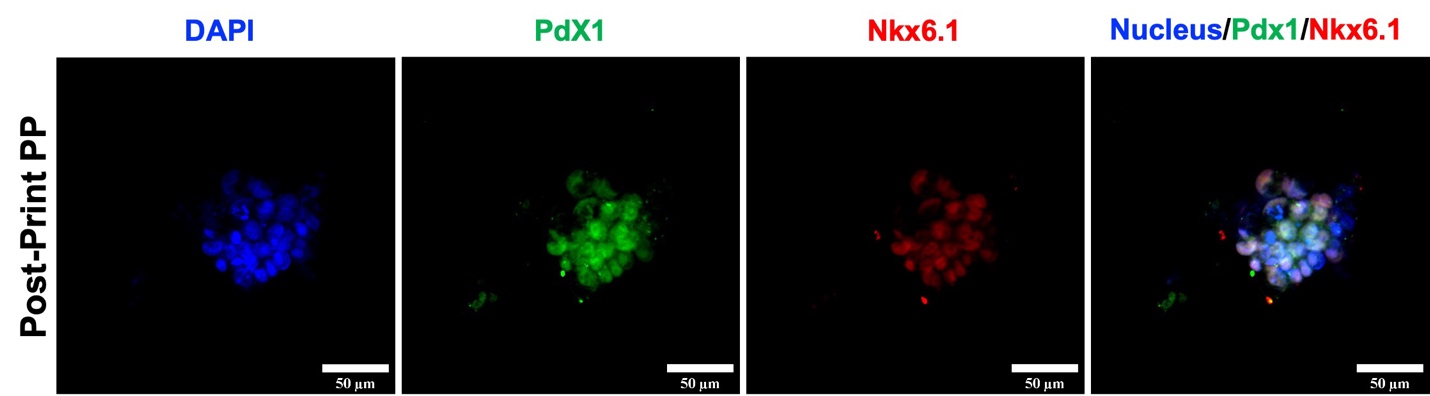

**Figure S12:** Undifferentiated iPSC-2 aggregates were printed and differentiated over 14 days to the pancreatic progenitor stage. Fluorescent staining images were taken 14 days after printing, where aggregates were fluorescently stained for nuclei (DAPI), pancreatic progenitor marker PDX1 (shown in green), and pancreatic progenitor marker NKX6.1 (shown in red). Scale bar representing 50 μm.

**Pre-Print PP**

**Day 1**

**Day 3**

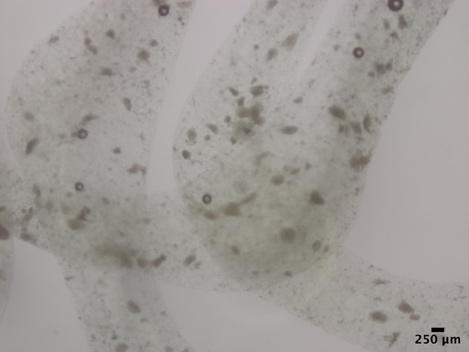

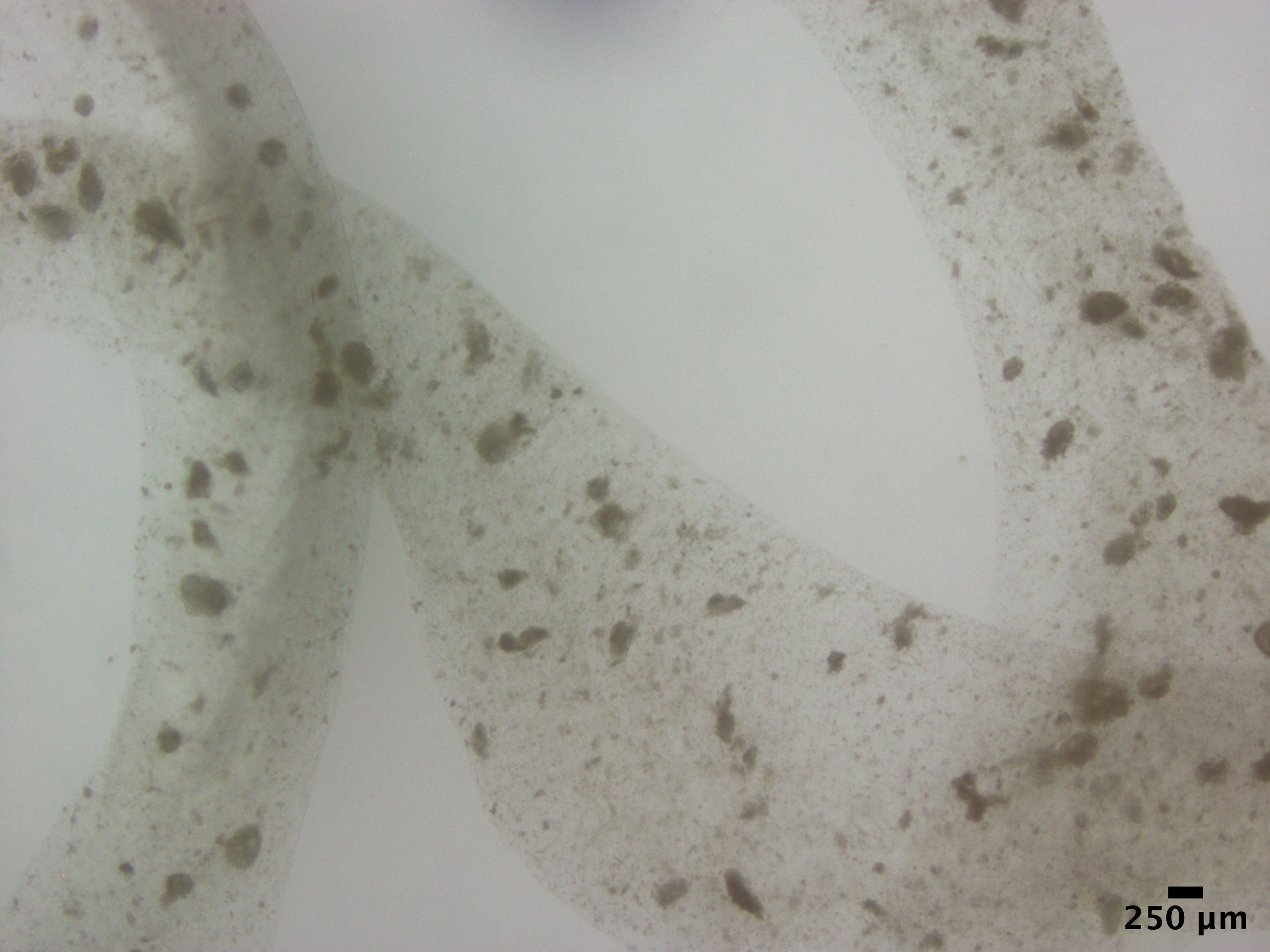

**Figure S13:** For pre-print PP (pancreatic progenitor), pancreatic progenitors iPSC aggregates were printed and maintained in culture for 3 days. Phase imaging for pre-print PP was taken on Days 1 and 3 post-printing. Scale bar representing 250 μm.

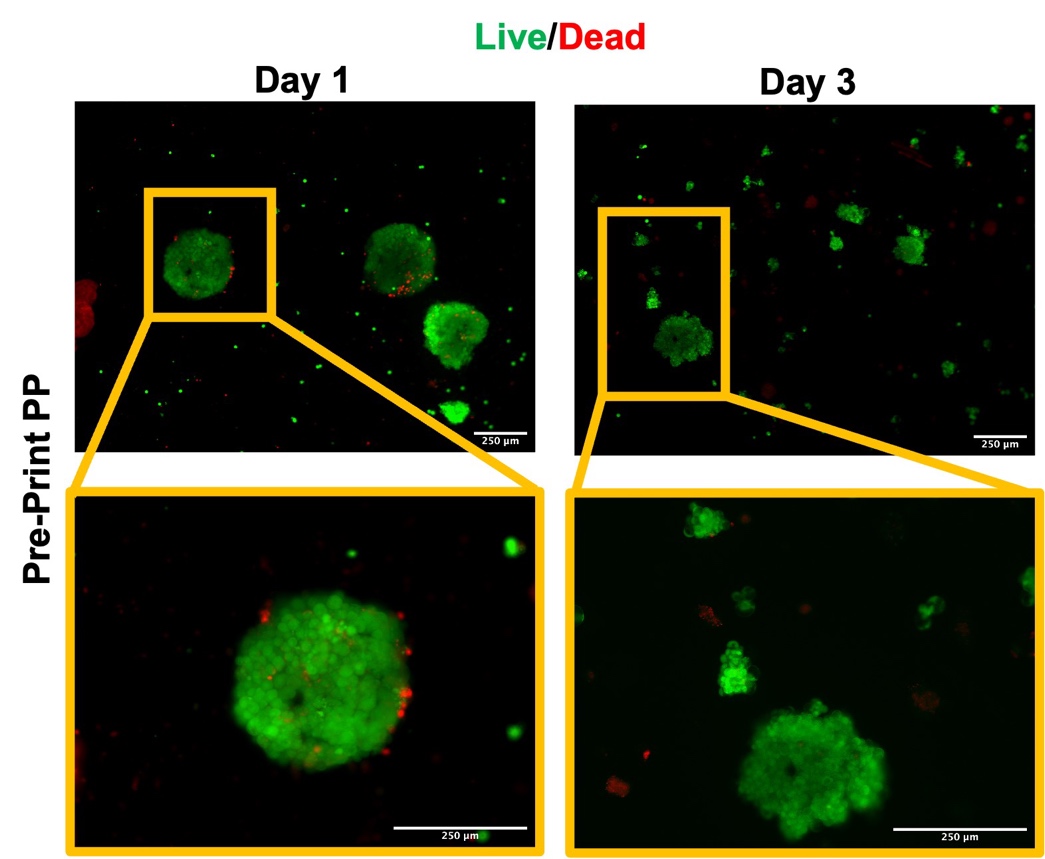

**Figure S14:** Live/Dead staining of printed iPSC pancreatic progenitor aggregates that were maintained in culture for 3 days post-printing. Aggregates were printed in a 3% w/v alginate/ 6% w/v methylcellulose bioink at 27 kPa and 12 mm s^-1^. All scale bars convey 100 μm. Live cells are shown in green, dead cells are shown in red.

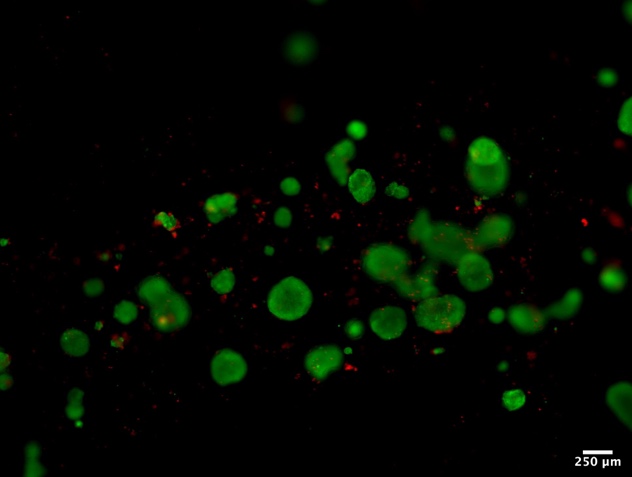

**Figure S15.** Representative viability staining for iPSC aggregates printed and still embedded within the construct. Live cells are shown in green, dead cells are shown in red. Scale bar conveys 250 μm.

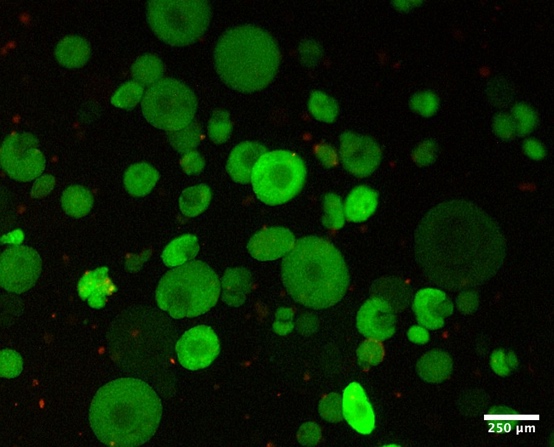

**Figure S16.** Representative LiveDead imaging of decapsulated iPSC derived mature islets, previously cultured in 1.5% w/v alginate. Live cells are shown in green, dead cells are shown in red. Scale bar represents 250 μm.

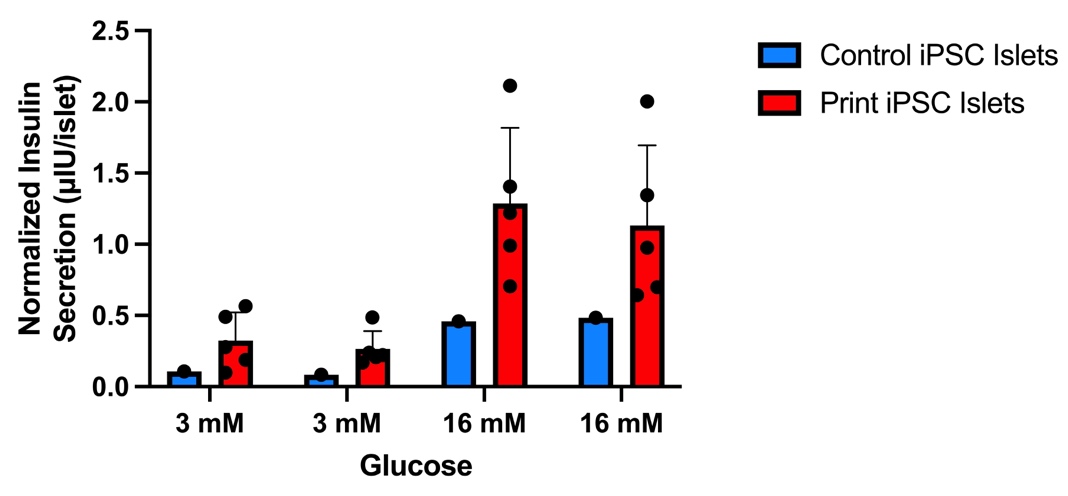

**Figure S17.** Glucose stimulated insulin secretion of printed and control iPSC-derived islets after 7 days in culture. Results for control iPSC-derived islets are shown in blue, and results for printed iPSC-derived islets are shown in red.

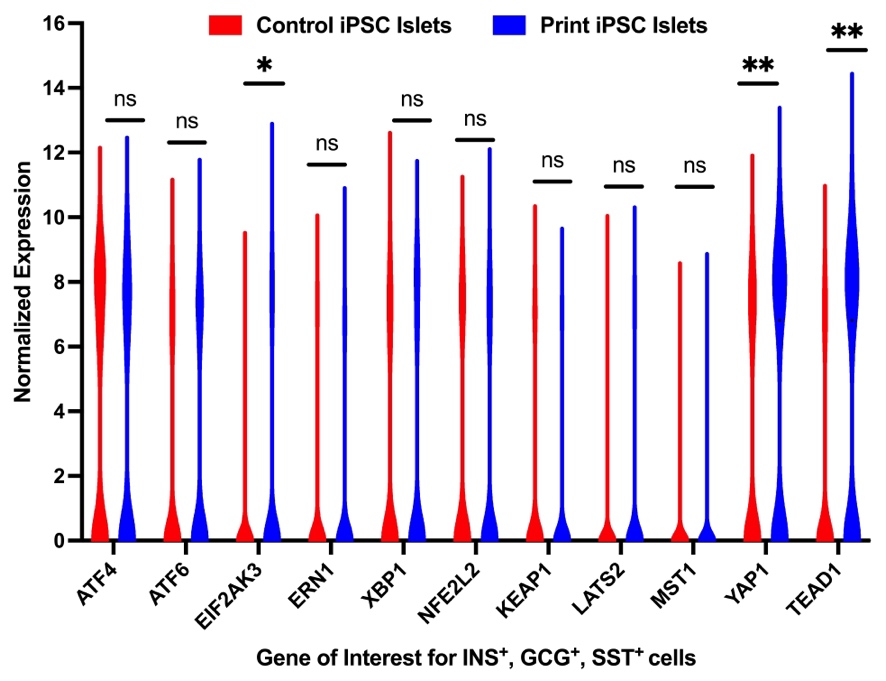

**Figure S18.** All endocrine cells were extracted from the general cell population in both the control and printed iPSC islets and examined for gene expression of key markers in the HIPPO, ER stress, and oxidative stress pathways. Violin plots were generated for the expression of ER Stress pathway genes of interest: *ATF4, ATF6, ERN1 (IREα), EIF2AK3 (PERK), XBP1,* Oxidative Stress pathway genes of interest: *NFE2L2 (NRF-2), KEAP1*, and HIPPO Signaling pathway: *LATS2, MST1, YAP1*, and *TEAD1*, indicating the level of expression (normalized counts) and the density of expression within the sample (ns: non-significant, *: p < 0.05, **: p < 0.01). Significant downregulation of expression was only noted in *EIF2AK3,* and for the HIPPO signaling pathway, *YAP1* and *TEAD1* were significantly upregulated in the printed sample.
